# Wnt4a is indispensable for genital duct elongation but not for gonadal sex differentiation in the medaka *Oryzias latipes*

**DOI:** 10.1101/2023.05.26.542371

**Authors:** Akira Kanamori, Ryota Kitani, Atsuko Oota, Koudai Hirano, Taijun Myosho, Tohru Kobayashi, Kouichi Kawamura, Naoyuki Kato, Satoshi Ansai, Masato Kinoshita

**Author notes:** CORRESPONDING AUTHOR: AKIRA KANAMORI, Division of Biological Science, Graduate School of Science, Nagoya University, Furocho Chikusa, Nagoya 464-8602, Japan. Graduate School of Human Development and Environment, Kobe University, Hyogo 657-8501, Japan (Ryota Kitani).

## Abstract

In most vertebrates, oviducts are derived from Mullerian ducts and sperm ducts from Wolffian ducts. In teleosts, however, Mullerian ducts are absent. Wolffian ducts function as nephric ducts in both sexes, and genital ducts are formed by posterior extension of either ovaries or testes. Whether genital ducts of teleosts are newly evolved organs or are a variant of the Mullerian ducts is an important question for evolutionary mechanisms of morphogenesis. One of the genes essential for Mullerian duct formation in mice, *wnt4*, is expressed in the mesenchyme and induces invagination of the coelomic epithelium and its posterior elongation. Here, we approached to the above question by examining genital duct development in mutants of two Wnt4 genes in medaka (*wnt4a* is orthologous to mouse *wnt4* and *wnt4b* paralogous). The *wnt4b* mutants had short body but were fertile with normal genital ducts. In contrast, both male and female *wnt4a* mutants had posterior elongation of the gonads stopped within or just outside the coelom, depending on the mutated alleles. Genetic females of the *scl* mutants (unable to synthesize testosterone or estrogens) have gonads containing both previtellogenic oocytes and spermatogenic cysts. Their gonads do not have ovarian cavities or sperm duct primordia and are lacking genital ducts completely. The results suggest Wnt4a target organs are posterior parts of the ovarian cavities or the sperm duct primordia. Medaka *wnt4a* was expressed in the mesenchyme ventral to the genital ducts in both sexes. Thus, the aborted elongation of genital ducts in the *wnt4a* mutants, the ortholog of mouse *wnt4*, suggests strongly that mouse Mullerian ducts and teleost genital ducts share homologous developmental processes. To further demonstrate this possible homology, mechanisms of genital duct formation and possible roles of Wnt4 should be compared before and after the appearance of Mullerian ducts in vertebrate phylogeny, namely jawless fish and cartilaginous fish. Additionally, *wnt4a* and *wnt4b* single mutants or double mutants did not show sex-reversal, suggesting both genes are dispensable for gonadal sex differentiation in medaka. This is in contrast to indispensable function of Wnt4 in mammalian ovarian differentiation.

## 1. INTRODUCTION

In most jawed vertebrates, oviducts and sperm ducts are derived from Mullerian ducts (MDs) and Wolffian ducts (WDs), respectively (see Romer and Parsons, 1977; Blüm, 1986; Lombardi, 1998). However, in teleosts, both male and female genital ducts are formed by posterior elongation of the gonads. Anatomically, there is no MDs in teleosts and WDs (mesonephric ducts) function as nephric ducts in both sexes throughout their life cycle. Therefore, these genital ducts in teleosts have been claimed by many researchers to be completely different and non-homologous organs to oviducts or sperm ducts of other jawed vertebrates (see above and Nagahama, 1983). We are interested in whether these genital ducts in teleosts are formed independently from MDs or share some developmental processes with MDs.

Developmental processes of the genital duct formation in teleosts have not been described in detail except in Japanese medaka, *Oryzias latipes* (Suzuki and Shibata, 2004). In males, mature sperm are liberated in canal-like spaces within the testes; posteriorly these canals fuse together to from single sperm duct primordium (here we call a central canal), which elongates posteriorly, leaves the coelom, elongates further beneath the urinary bladder, and finally fuse to the urethra from its ventral side; therefore, medaka has a single urogenital opening (see Fig. 1 and Fig. 8A for a diagram). In females, the process is rather complex. First, the ovarian cavity is formed by joining lateral elongation of tissue sheets from the dorso-central somatic cells with dorso-central elongation of the most lateral periphery of the somatic tissue sheets. Ovarian lamella containing developing oocytes and oogonia are thereby located ventrally to the ovarian cavity (Kanamori et al., 1985; Suzuki and Shibata, 2004; see Fig. 3 and Fig. 8D for a diagram). The most posterior portions of the ovarian cavity are devoid of the germ cells. The sphincter muscle beneath the urinary bladder protrude anteriorly into the coelom from its ventral side and envelope lumen of the ovarian cavity to form oviduct primordium, which then elongate posteriorly within the sphincter muscle, finally open at the dorsal base of the urogenital papillae (UGP) well developed in the females.

**Figure 1.**
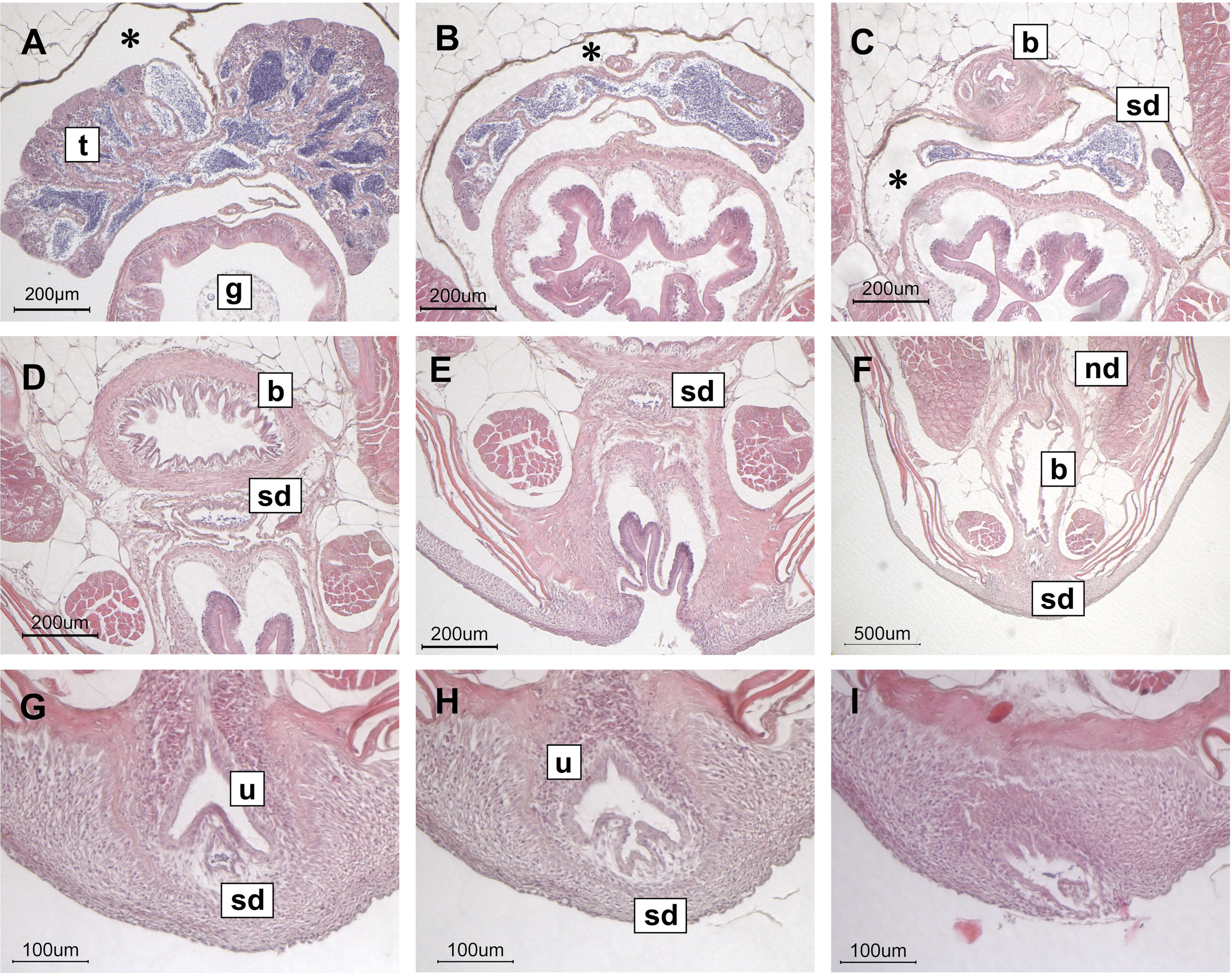
Histology of a wildtype male (G_6_ from incrossing G_5_ heterozygotes) at 4 months after hatching. Cross sections, A to I, are shown in anterior to posterior order. In medaka, single testis (t) is situated centrally in the coelom above the gut (g) (A). Mature sperm were liberated from the spermatogenic cysts and were present in the canal like spaces (dark stained lumens in A), which posteriorly fuse together to form larger lumens (B). Further posteriorly, the lumens fused to make a single central canal (sperm duct, sd); no spermatogenic cysts were observed at this level (C). Just above the coelom, anterior end of the urinary bladder (b) was shown. The sperm duct left the coelom beneath the urinary bladder (D). The sperm duct was shown between the anus and the urinary bladder (E). The nephric ducts (nd) were soon to join the urinary bladder (F). The sperm ducts finally fused the urethra (u) from its ventral side (G and H) and the urinogenital duct opened (I). A sideview diagram is shown in Fig. 8A. See also movie S1 for gross anatomy of the urogenital system of the male medaka.

Cellular and molecular processes of the MD development are most well documented in mice, where 20-30 genes have been identifies as indispensable in MD formation (see Mullen and Behringer, 2014). Interestingly, most of these genes are expressed in both MD and WD tissues.

One of the MD specifically expressed gene is *wnt4*, a member of Wnt ligand family known to function in various aspects in development (see Logan and Nusse 2004; Steinhart and Angers, 2018; Mehta et al., 2021), expressed in the mesenchyme to induce invagination of the coelomic epithelium and posterior elongation of the MDs along the WDs. Therefore, in this study, we made mutants of the medaka *wnt4* homologs (*wnt4a* and *wnt4b*) and analyzed their genital duct development. No apparent phenotype may suggest completely independent origin of the teleost genital ducts from MDs and any abnormal development in the mutants may indicate that development of the teleost genital ducts share some homologous developmental processes to those of the MDs in jawed vertebrates.

## 2. MATERIALS AND METHODS

### 2.1. Medaka

Inbred strains of the Japanese medaka (*Oryzias latipes*), HdrR, established in National Institute of Radiological Sciences, Japan (Hyodo-Taguchi and Sakaizumi, 1993), were used for *wnt4a Δ*30 mutants, and Cab, established by Carolina Biological Supply Company (Loosli et al., 2000), were used for all other mutants. Adults, embryos, and hatchlings were kept at 26–28 °C with 14L:10D light cycle. The Animal Care and Use Committee of Nagoya University approved all husbandry and experimental procedures in the present study.

### 2.2. *wnt4a* and *wnt4b* editing by CRISPR/Cas9

Target sites were selected with a web tool detecting micro-homology sites (http://viewer.shigen.info/cgi-bin/crispr/crispr.cgi) (see Fig. S2). Short guide RNAs to the target sites and a Cas9 mRNA were co-injected into fertilized eggs according to Ansai and Kinoshita (2014). Homologous recombination were induced as described in Murakami et al., 2017. Target sites, mutated alleles, and expected proteins are described in Fig. S2. Injected individuals were crossed with wildtype and F_1_ were used for identifying mutated alleles. The F_1_ with mutated alleles were then backcrossed at least once before incrossing heterozygotes (here, the first progeny between heterozygotes is named G_1_) and the mutants were maintained as heterozygotes. The most of the mutants can be obtained through National Bioresource Project Medaka (https://shigen.nig.ac.jp/medaka/).

### 2.3. Genotyping

Genomic DNA for genotyping was extracted from embryos, tail fin clips, or head tissue by alkaline lysis methods described in Ansai and Kinoshita (2014). PCR was performed with ExTaq (TaKaRa, Japan) on 2720 thermal cycler (Applied Biosystems). Amplified products were analyzed with conventional agarose gels, followed with a microchip electrophoresis analyzer, MultiNA, with DNA-500 reagents (Shimadzu, Japan). With the latter, hetero-duplexed amplified DNA from wild type and mutated alleles can be visualized easily. In medaka, sex is determined genetically with XX/XY system, where only Y chromosome contains male determining gene, *dmy*. For determining genetic sex, two primer pairs, 17. 19 /17. 20 and 17. z1 /17. z2 were used (Matsuda et al., 2002; Fig. S2). *wnt4a* and *wnt4b* mutant alleles were genotyped as described in Fig. S2. PCR amplified products from mutated alleles were sequenced with Big Dye terminator (v3.1) and Prism 3100 Genetic Analyzer (Applied Biosystems). *Scl* mutants were genotyped as described in Sato et al. (2008).

### 2.4. Morphological analyses

Histological slides were prepared in conventional methods from samples fixed with Bouin’s, embedded in paraffin (Sigma-Aldrich, USA), sectioned at 7 μm, and stained with hematoxylin and eosin (Muto Pure Chemicals, Japan).

### 2.5. *In situ* hybridization

In situ hybridization was performed as described previously (Horie et al., 2016). Briefly, 7 μm paraffin sections were hydrated, treated with 1 μg/ml proteinase K (Sigma-Aldrich, USA) for 10 min at 37 ℃, and hybridized with DIG-labeled antisense RNA probes. Detection was done with alkaline phosphatase-labeled anti-DIG antibody and NBT/BCIP (Roche diagnostics, USA). Control experiments with sense probes did not give detectable signals (data not shown). A near full-length cDNA of *wnt4a* (nucleotide 106-1018, GenBank# NM_001160439) was used for a probe.

## 3. RESULTS

### 3.1. *wnt4b* mutants

*Wnt4* is conserved in almost all animal groups (Fig. S1). As already shown by Kossack et al. (2019), jawed vertebrates have two Wnt4 genes: *wnt4a* and *wnt4b*. Some groups, including mammals, lack *wnt4b*. Mammalian *wnt4* is orthologous to medaka *wnt4a* and paralogous to medaka *wnt4b* (Fig. S1). The phenotypes of a natural *wnt4b* mutant (little expression of *wnt4b*, presumably by a transposon insertion in its promoter) have been described by Inohaya et al. (2010). Similar to the natural mutant, our induced mutants (homozygous *Δ*5 and *Δ*29 mutants, encoding only a part of the signal peptide; Fig. S2) had extremely shortened body (Fig. S3). But they were fertile and histological examination showed proper development of the genital ducts in both males and females (data not shown).

### 3.2. *wnt4a Δ*30 mutant

We first analyzed a *wnt4a* mutant *Δ*30, which has a 30 base deletion overlapping the initiation codon (Fig. S2). The deletion would move translation initiation to the second AUG, deleting first 11 residues of the 22 signal peptides. Generally, signal peptides for secretion consist of N-, H-, and C-regions, all of which are important for proper secretion (see Izard and Kendall,1994; Owji et al., 2018). The *wnt4a Δ*30 allele lack entire N-region containing positive charged residues. In fact, *wnt4a Δ*30 allele encoded protein is not recognized as a secretion signal by a web-based prediction program (https://services.healthtech.dtu.dk/services/SignalP-6.0/). At G_1_, G_2_, and G_6_, we histologically analyzed adults of both sexes (3-6 months after hatching). The wildtype and heterozygotes males had normal sperm ducts (Fig. 1). Medaka have a single gonad suspended in the coelom dorsal to the gut. In contrast, homozygous mutant males had normal testes anteriorly (Fig. 2A and B) and possessed posterior extension of the gonad devoid of germ cells (sperm duct primordium, Fig. 2C and D) but they were closed within the coelom (Fig. 2E and F). No sperm ducts were observed outside the coelom. In females also, the wildtype and heterozygotes had normal oviducts (Fig. 3). In contrast, as in males, homozygous mutant females had normal ovaries with ovarian cavities anteriorly (Fig. 4A) and possessed posterior extension of the ovarian cavities devoid of germ cells (oviduct primordia, Fig. 4B and C) but they were closed within the coelom (Fig. 4D). No oviducts were observed outside the coelom (Fig. 4E) and the muscular tissue was present beneath the urinary bladder. In some females, the muscular tissue was fully developed and protruded anteriorly into the coelom; posterior end of the ovarian cavity was enveloped by the muscle tissue within the coelom (Fig. S4). The cumulative results of both sexes are summarized in Table 1. We found one mutant male having normal sperm duct and two heterozygous females having oviducts that were enveloped within the muscle and closed outside the coelom beneath the urinary bladder (diagrammed in Fig. 8I). These two females had undeveloped UGP, suggesting they were immature females. Sideview diagrams (Fig. 8) are presented for better understanding of anatomy of medaka urogenital systems.

**Figure 2.**
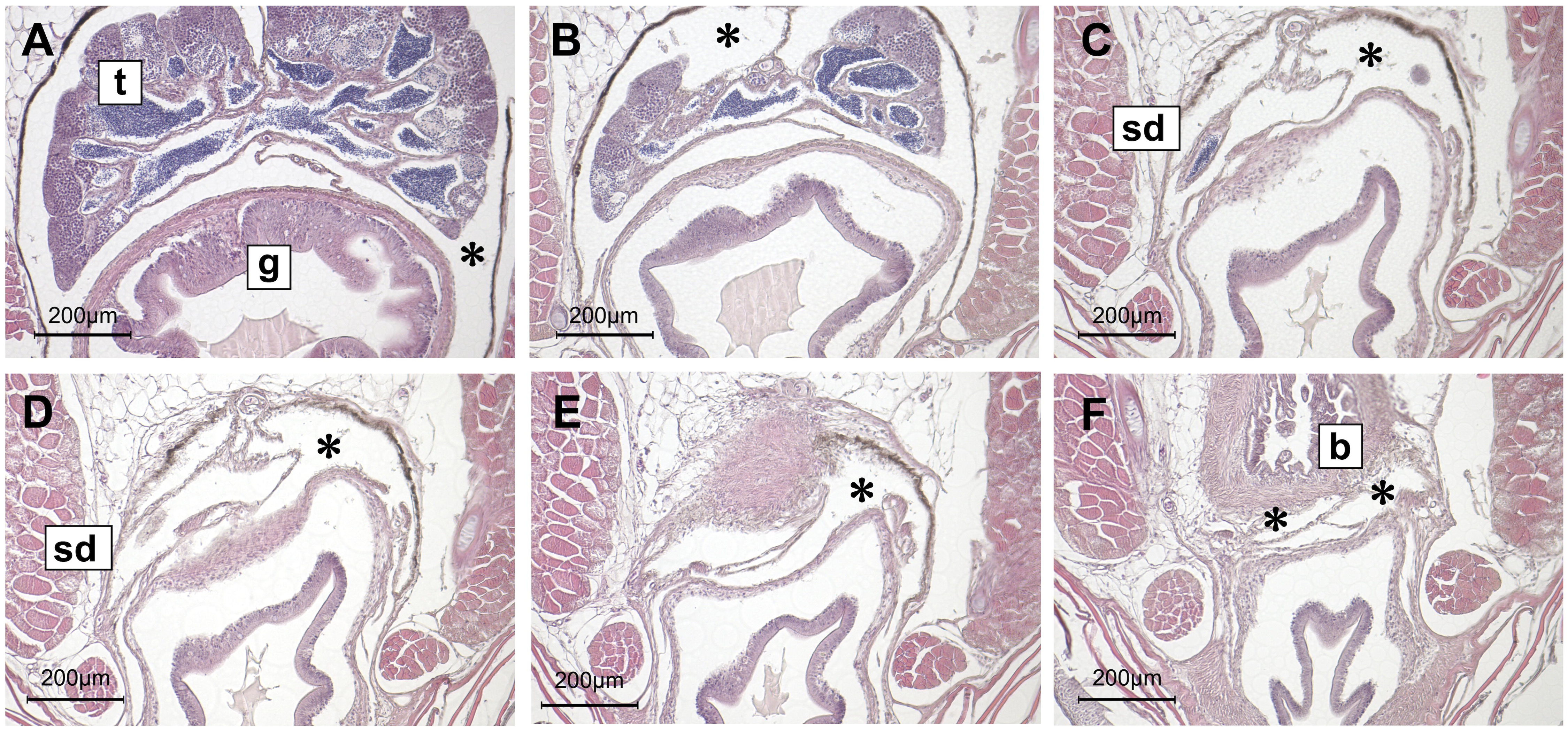
Histology of a male homozygous for *wnt4aΔ*30 (G_6_ from incrossing G_5_ heterozygotes) at 4 months after hatching. Cross sections, A to F, are shown in anterior to posterior order. Spermatogenesis seemed progressing normally (A) and a sperm duct primordium (central canal) was formed posteriorly (B and C). However, the primordium stopped elongating within the coelom (D, E, and F). t, testis; g, gut; sd, sperm duct (primordium); b, urinary bladder; asterisks, coelom. A sideview diagram is shown in Fig. 8B.

**Figure 3.**
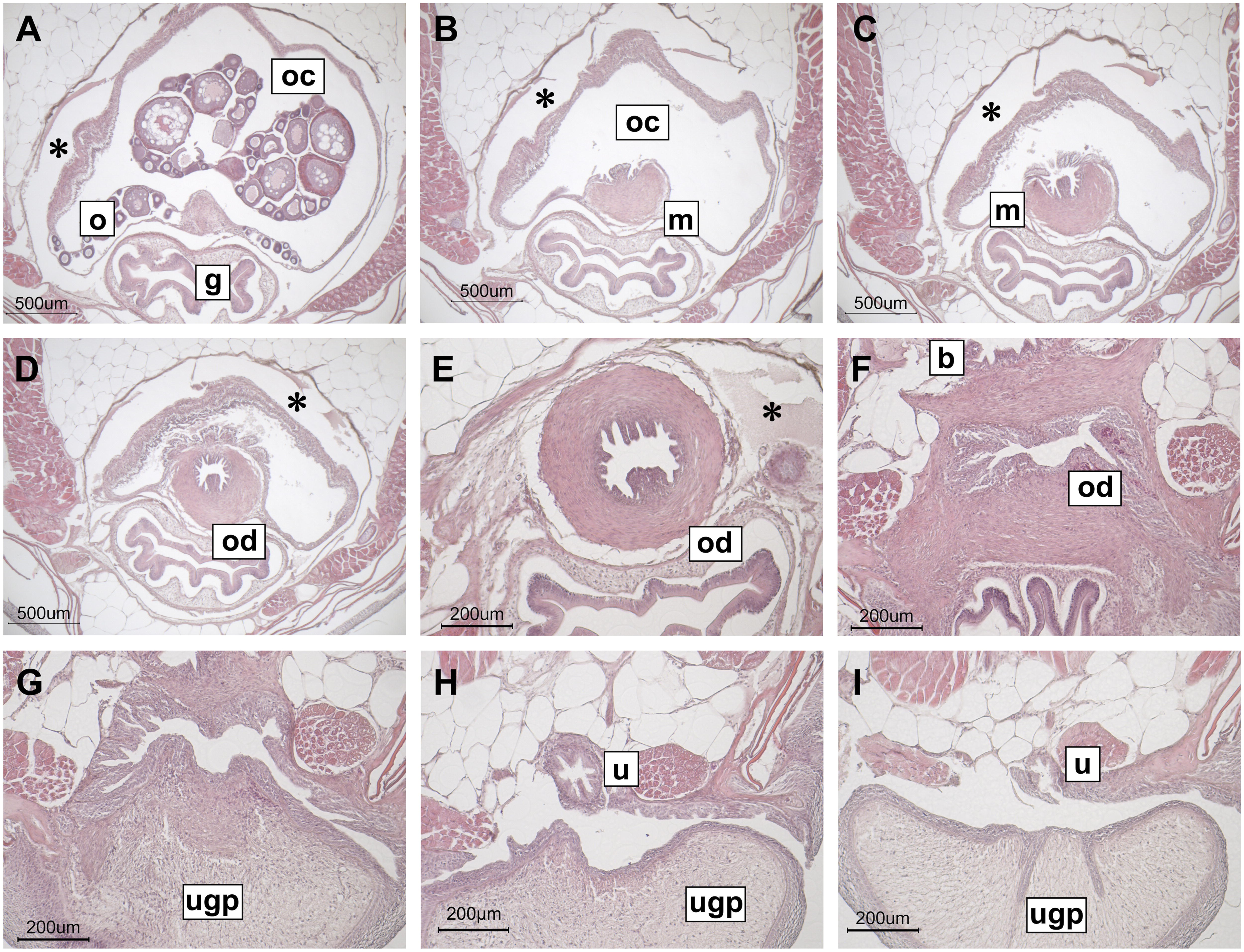
Histology of a wildtype female (G_6_ from incrossing G_5_ heterozygotes) at 4 months after hatching. Cross sections, A to I, are shown in anterior to posterior order. In medaka, single ovary (o) is situated centrally in the coelom (asteriks) above the gut (g) (A). The ovarian lamella containing oogonia and developing oocytes were shown at the ventral side and the center of the ovarian cavity (oc). The posterior part of the ovarian cavity (oviduct primordium) did not contain germ cells and the sphincter muscle (m) protruded from posterior was shown (B). The part of the ovarian cavity was enveloped by the sphincter muscle (C) to make the oviduct (od) posteriorly (D and E). The oviduct then left the coelom, elongated under the urinary bladder (b) (F), and opened at the dorsal side of the base of the urogenital papillae (ugp), which was developed fully (G). The urethra (u) opened separately from the oviduct between body and the ugp (I). A sideview diagram is shown in Fig. 8D. See also movie S2 for gross anatomy of the urogenital system of the male medaka.

**Figure 4.**
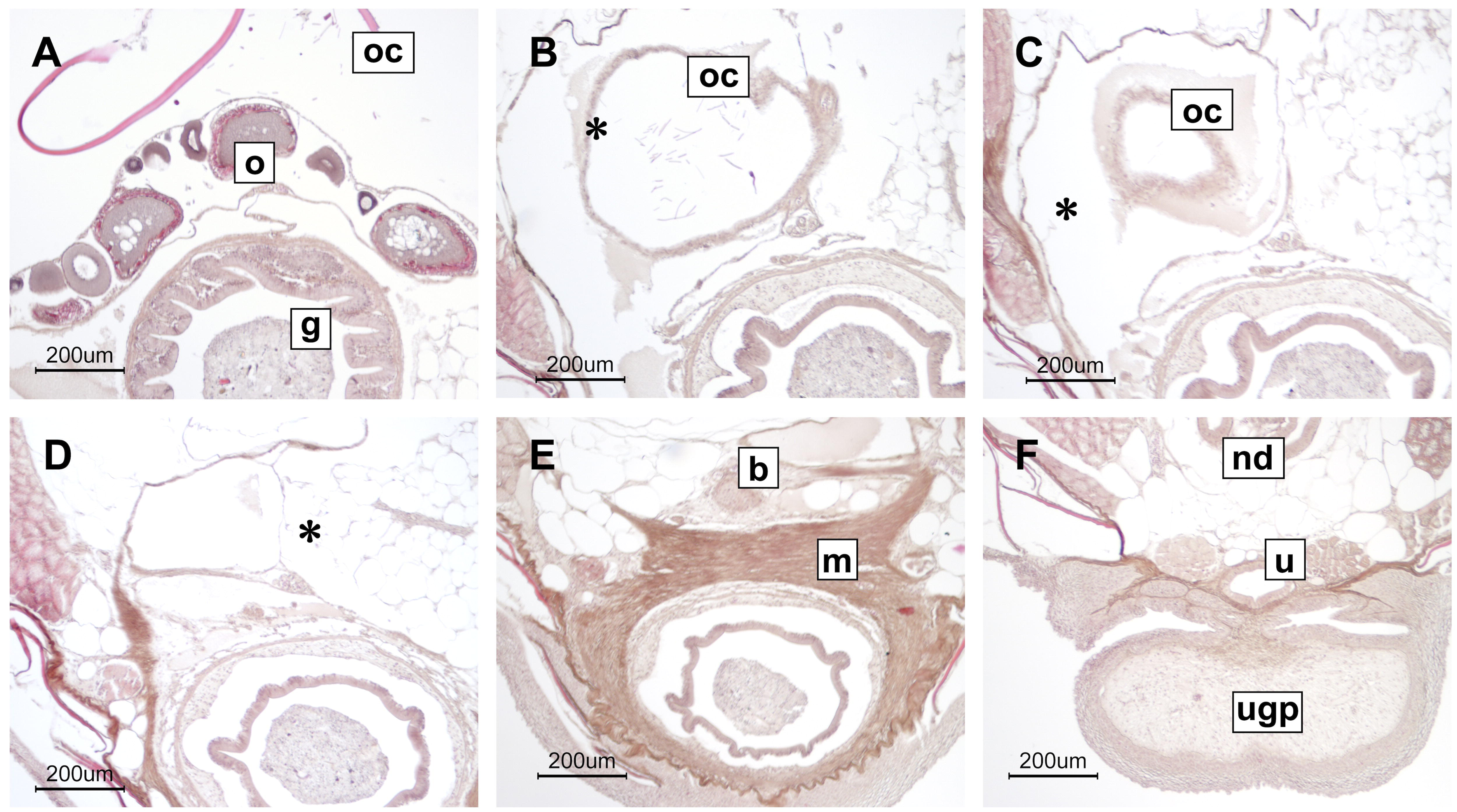
Histology of a female homozygous for *wnt4aΔ*30 (G_2_ from incrossing G_1_ heterozygotes) at 4 months after hatching. Cross sections, A to F, are shown in anterior to posterior order. Oogenesis seemed progressing normally (A) and a posterior portion of ovarian cavity (oc) devoid of germ cells (oviduct primordium) was formed (B). However, the primordium stopped elongation within the coelom (C and D). Muscular tissue beneath the urinary bladder, well developed urogenital papillae, and the urethra were developed normally. o, ovary; g, gut; b, urinary bladder; ugp, urogenital papillae; nd, nephric duct; asterisks, coelom. A sideview diagram is shown in Fig. 8E.

**Table 1.**
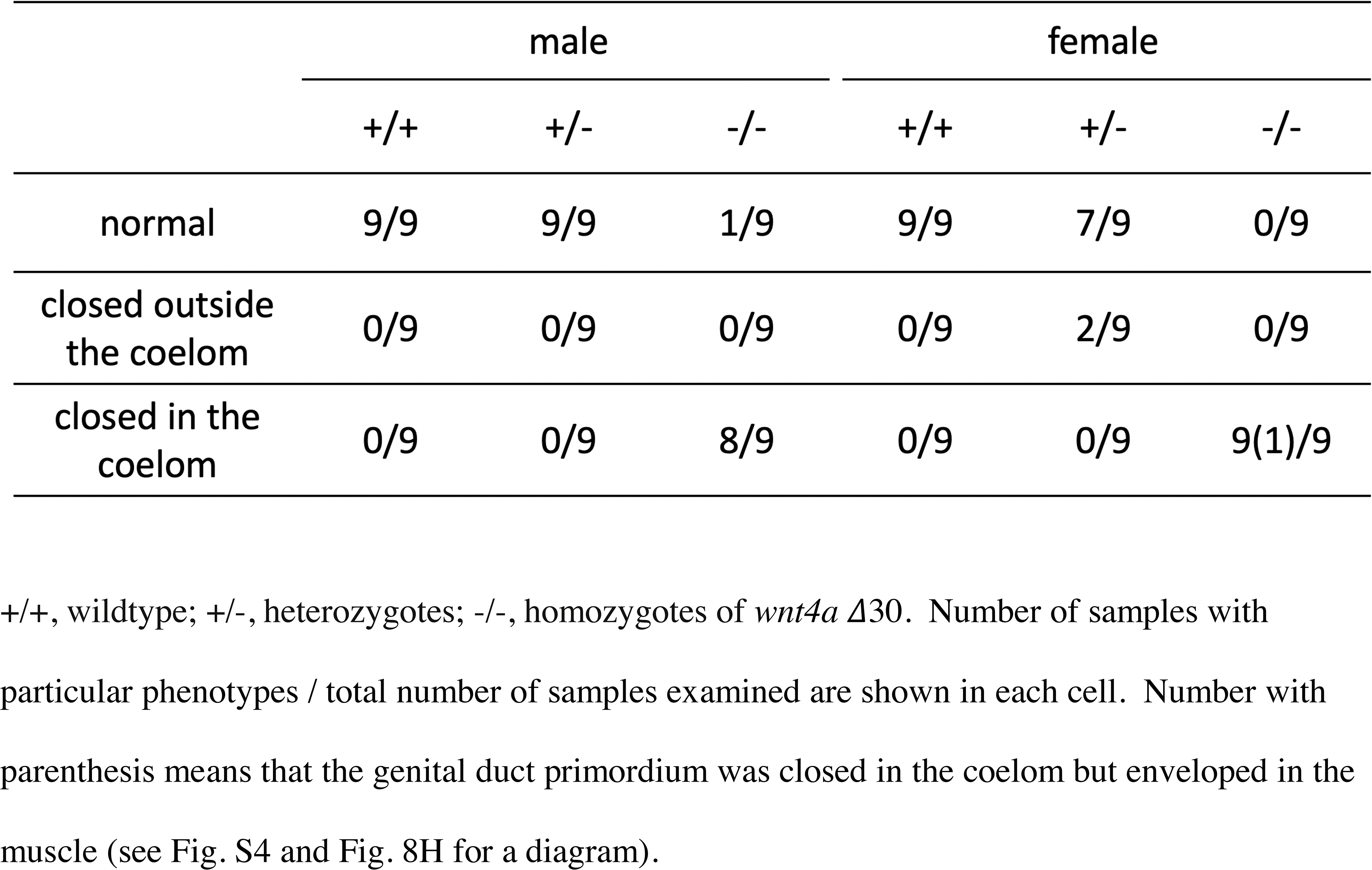
Genital ducts phenotypes of *wnt4a Δ*30 progenies (G_1_, G_2_, and G_6_ combined) +/+, wildtype; +/-, heterozygotes; -/-, homozygotes of *wnt4a Δ*30. Number of samples with particular phenotypes / total number of samples examined are shown in each cell. Number with parenthesis means that the genital duct primordium was closed in the coelom but enveloped in the muscle (see Fig. S4 and Fig. 8H for a diagram).

### 3.3. Other *wnt4a* mutated alleles

As noted above, the *Δ*30 allele of *wnt4a* may not be a null allele. Therefore, we examined other mutated alleles induced with CRISPR/Cas9 mediated homologous recombination (see Fig. S2). One knock-in allele has an insertion containing mouse crystalline promoter followed by GFP or mCherry and SV40 polyA signal (Pcry-GFP or Pcry-mCherry) in the third exon. The fluorescent proteins are expressed in the lenses so that genotyping becomes easier. Another knock-in alleles were made with GFP or mCherry fused in-frame with *wnt4a* coding sequence in the third exon (GFP-fusion or mCherry-fusion). Interestingly, these knock-in alleles showed less severe phenotypes than those of the *Δ*30 allele. Heterozygotes of GFP-fusion were crossed with those with mCherry-fusion and adults of the progenies were histologically examined at G_1_ and G_2_. Five of 8 male heterozygotes with both GFP-fusion and mCherry-fusion had sperm ducts closed outside the coelom (Fig. 5A-D; table 2). The posterior sperm duct primordium elongated out of the coelom but closed beneath the urinary bladder (Fig. 5C and D). The remaining 3 mutants had normal sperm ducts joining urethra. In contrast, all but one female mutants with GFP-fusion and mCherry-fusion had closed oviduct primordium within the coelom. The wildtype and the heterozygotes had normal genital ducts. The Pcry mutants had least severe phenotypes (Fig. 5E-H; table 3). All male mutants of the Pcry-GFP homozygotes and the Pcry-mCherry homozygotes and 2/3 of the Pcry-GFP / Pcry- mCherry heterozygotes had normal sperm ducts joining urethra. Only one of the Pcry-GFP / Pcry- mCherry heterozygotes had sperm duct closed out of the coelom. The oviduct phenotypes of the female mutants (Pcry-GFP homozygotes, Pcry-mCherry homozygotes, Pcry-GFP / Pcry-mCherry heterozygotes) were divided about equally into normal, closed outside the coelom, closed within the coelom. The second phenotype were only observed in the Pcry mutants; the posterior elongation of the ovarian cavity apparently left the coelom (Fig. 5G) but closed within the muscle. All of the heterozygotes had normal genital ducts. I addition to the knock-in alleles, we found out deletion alleles (*Δ*8 and *Δ*14 in the third exon; Fig. S2) from F_1_ of the GFP-fusion experiments. We could not obtain more than 3 male mutants but both male and female mutants of the heterozygotes showed closed genital duct phenotypes less severe than the *Δ*30 mutants (table S1).

**Figure 5.**
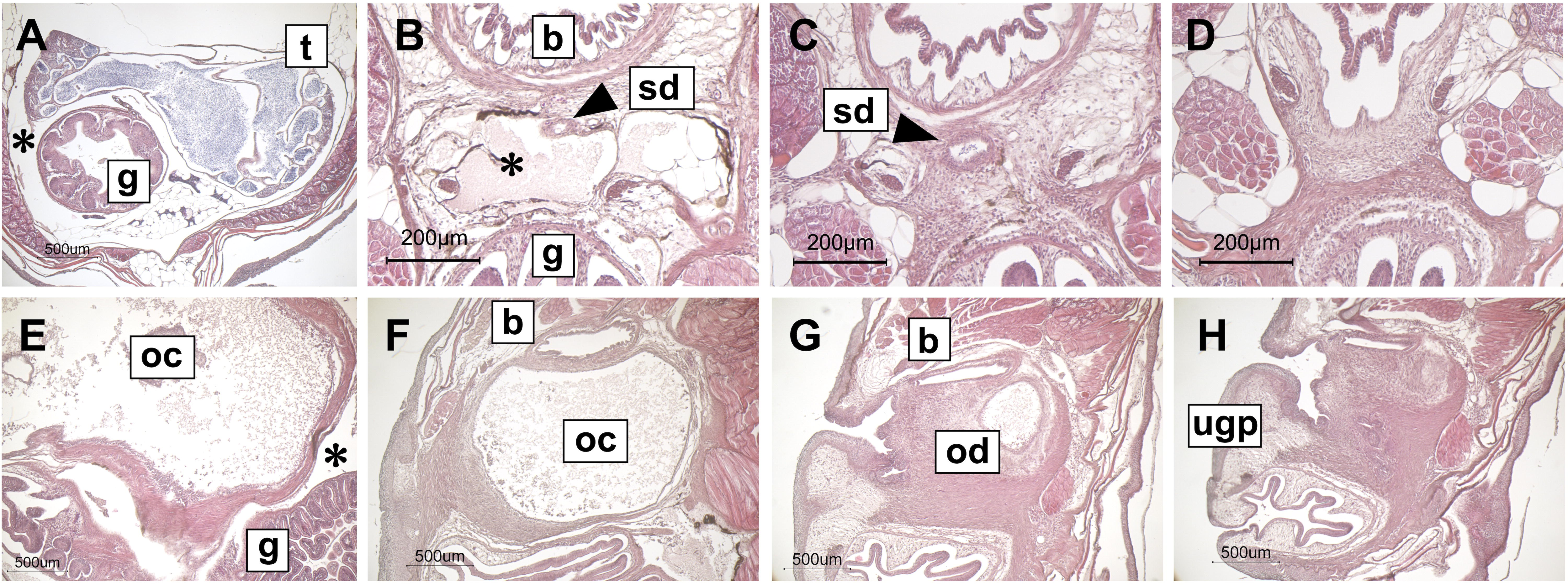
Some *wnt4a* mutants from knock-in alleles showed termination of the genital ducts outside the coelom. Cross sections, A to D for a male heterozygous for GFP-fusion/mCherry-fusion and E-H for a female homozygous for Pcry-GFP, are shown in anterior to posterior order. The sperm duct (sd, arrowheads) or oviduct (od) elongated outside the coelom and stopped beneath the urinary bladder (C and D for the male; G and H for the female). t, testis; g, gut; b, urinary bladder; sd, sperm duct; oc, ovarian cavity; g, gut; b, urinary bladder; ugp, urogenital papillae; asterisks, coelom. Sideview diagrams are shown in Fig. 8C and 8F.

**Table 2.**
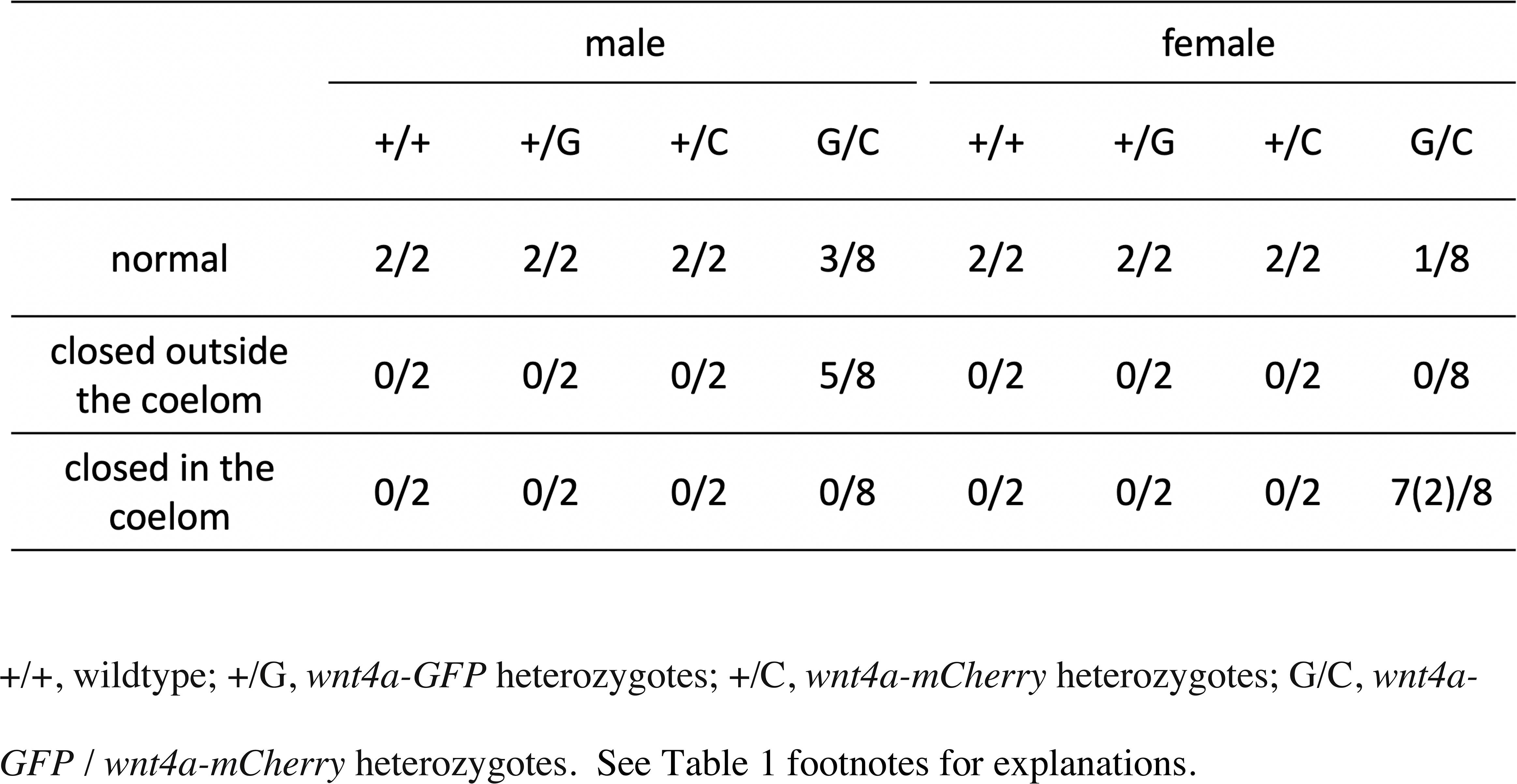
Genital ducts phenotypes of *wnt4a* fusion allele progenies (G_2_) +/+, wildtype; +/G, *wnt4a-GFP* heterozygotes; +/C, *wnt4a-mCherry* heterozygotes; G/C, *wnt4a- GFP* / *wnt4a-mCherry* heterozygotes. See Table 1 footnotes for explanations.

### 3.4. Genital ducts of *scl* mutant medaka

As shown in the previous sections, loss of function of *wnt4a* leads to premature termination of posterior elongation of the genital ducts. However, the mutants did have duct primordia: posterior central canals in the males and posterior ovarian cavities in the females. These are most likely target tissues responsive to Wnt4a. The *scl* mutant medaka do not produce androgens and estrogens because of a *17, 20-lyase* mutation (Sato et al., 2008). Total of 7 adult mutants (5 months after hatching) were histologically examined. Two genetic males (i.e. XY) and one out of 5 genetic females (XX sex-reversed) had normal testes with normal sperm ducts joining urethra (data not shown). Four remaining XX had ovary-like gonads containing previtellogenic oocytes together with scattered spermatogenic cysts (Fig. 6 A-D). The gonads did not possess ovarian cavities as already described in Sato et al., (2008). The formation of the ovarian cavity depends on estrogens (Suzuki et al., 2004). The gonads contained some spermatogenic cysts containing spermatids but no canals for sperm were present. The most posterior part of the gonads contained only somatic cells without lumens and no further elongation were observed within the coelom (Fig. 6E-H).

**Figure 6.**
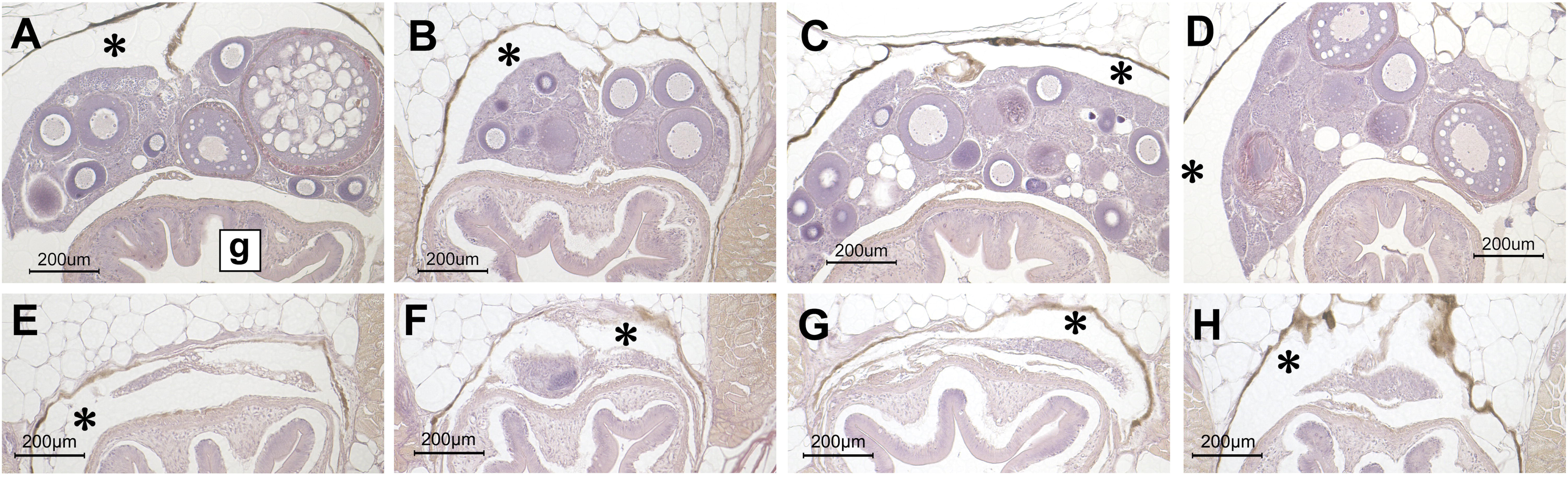
Histology of gonads from 4 XX adults homozygous for *scl* at 5 months after hatching. Cross sections of the posterior part of the gonads, showing ovary-like structure containing previtellogenic oocytes with scattered spermatogenic cysts; no ovarian cavities or canals for sperm were formed (A to D). The posterior end of the gonads contained only somatic cells without lumens (E to H). A, B, C, and D and E, F, G, and H are from same individuals, respectively. g, gut; asterisks, coelom. A sideview diagram is shown in Fig. 8G.

These results further corroborate the hypothesis that duct primordia (posterior central canals in the males and posterior ovarian cavities in the females) are target tissue responsive to Wnt4a.

### 3.5. *wnt4a* expression during genital duct elongation

First, we tried immunohistochemical detection with GFP antibodies on medaka with the *wnt4a* GFP-fusion allele. However, we could not detect reasonable signals. Next, we used conventional *in situ* hybridization and detected signals on mesenchyme beneath the sperm ducts and epithelium of the sperm duct in the males (Fig. 7A) and mesenchyme beneath oviducts and periphery of the sphincter muscle in the females (Fig. 7B). In the gonads, spermatids and spermatozoa were stained because of their intrinsic alkaline phosphatase activities and the developing oocytes had positive signals; We did not detect positive signals on the gonadal somatic cells in both sexes (data not shown). The control sense probes did not give any detectable signals (data not shown).

**Figure 7.**
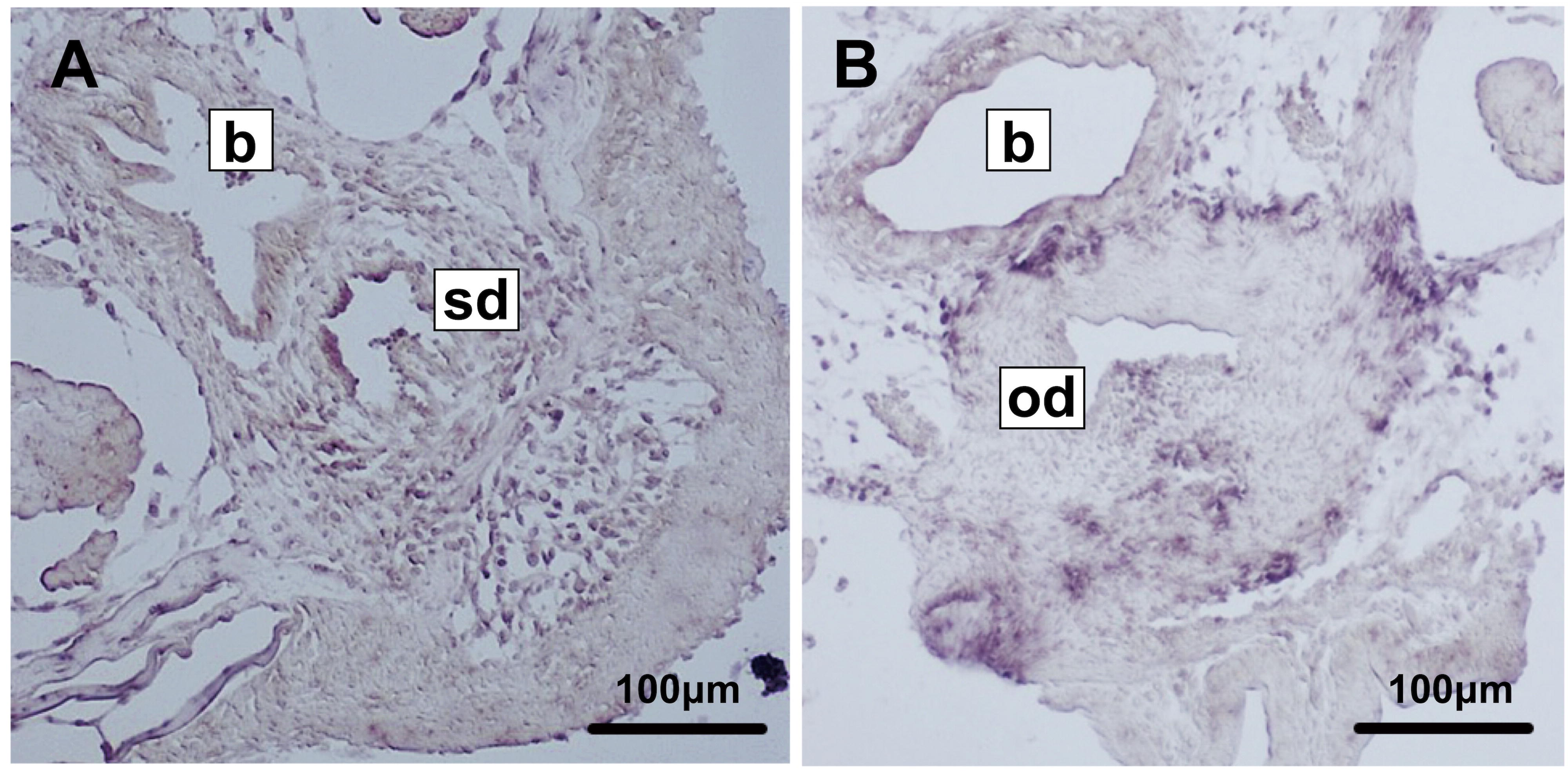
*In situ* hybridization of *wnt4a*. The cross sections of a male and a female at 70 days after hatching were hybridized to an antisense RNA probe. Signals were detected on mesenchyme beneath the sperm ducts and epithelium of the sperm duct in the male (A) and mesenchyme beneath oviducts and periphery of the sphincter muscle in the female (B). b, urinary bladder; sd, sperm duct; od, oviduct.

### 3.6. No sex-reversals in *wnt4* mutant medaka

The role of Wnt4 in ovarian differentiation is well known in mammals (Vainio et al., 1999; see also Bernard and Harley, 2007; Nicol and Yao, 2014); *wnt4* mutant females show partial sex-reversals. In teleosts, zebrafish *wnt4a* mutants showed higher male ratio than the wildtype (Kossack et al., 2019). Therefore, we examined whether sex-reversal occurs in *wnt4* medaka mutants. First, we checked 8 genetic males (XY) and 8 genetic females (XX) homozygous for *wnt4b Δ*29 allele and found no sex-reversals. Genetic sex was confirmed by the presence or absence of *dmy*, the male- determining gene of medaka with two independent primer pairs (Fig. S2). Phenotypic sex was determined by external morphology of dorsal and anal fins and UGP, together with a gross morphology of the gonads. For *wnt4a*, *Δ*30 and fusion alleles were examined and no significant sex-reversals were identified (Table S2). As *wnt4b* mutants are fertile, the G_6_ females homozygous for *wnt4b Δ*29 or *Δ*5 were crossed with the G_6_ male heterozygous for *wnt4a Δ*30. Progenies heterozygous for both *wnt4a* and *wnt4b* were incrossed to make *wnt4a* and *4b* double mutants. As *wnt4b* mutant phenotype (shorter body) can be detectable at a week after hatching, we only selected *wnt4b* homozygotes for further analyses. At adults, their genotypes and phenotypes were analyzed. As shown in table 4, no significant sex-reversals were identified in the double mutants. Genital duct phenotypes were further examined for some of those individuals. There seemed to be no additional effect of *wnt4b* mutation, either *Δ*29 or *Δ*5, on *wnt4a Δ*30 mutation (Table S3). We did not identify significant differences; most of the *wnt4a / wnt4b* double mutants had their genital ducts closed in the coelom similar to *wnt4a* single mutants (Table 1).

## 4. DISCUSSION

### 4.1. *wnt4a* but not *wnt4b* is indispensable for genital duct elongation in male and female medaka

The *wnt4a Δ*30 homozygous mutants of both sexes showed premature termination of genital duct elongation in the coelom (Fig. 1-4; Table1). The homozygous mutants of other *wnt4a* mutated alleles also showed similar but less severe phenotypes (Fig. 5; Table 2-4 and S1). Together, these results strongly indicate that in medaka, *wnt4a* is indispensable for genital duct elongation in both sexes. The similar anomalies of genital duct formation were reported in other teleost, zebrafish (Kossack et al., 2019). Male mutants had genital duct primordia but the left and right primordium did not fuse and elongate posteriorly. The female mutants also had undeveloped oviduct primordium, which did not elongate posteriorly. Expression of *wnt4a* was detected in mesenchyme beneath nephric ducts in both sexes in zebrafish (Kossack et al., 2019). Here, in medaka, we also detected *wnt4a* expression beneath the urinary bladder in both sexes. Taken together, both in medaka and zebrafish (two of the relatively diverged teleost species), *wnt4a* is expressed in the mesenchyme, suggesting hypothesis that Wnt4 directly or indirectly acts on the duct primordia to induce their posterior elongation. The *scl* mutants did not possess the duct primordia (posterior central canal in the males and posterior ovarian cavities in the females) and completely lacked genital ducts (Fig. 6), which corroborate further the above hypothesis. However, Wnt signals usually travel only several cell lengths (see Logan and Nusse 2004; Steinhart and Angers, 2018; Mehta et al., 2021). Here, the possible target tissue is located within the coelom. Therefore, how Wnt4a signals, directly or indirectly, acts on the possible target is unknown. Further cellular and molecular analyses of Wnt4a function in genital duct elongation should be performed in details.

### 4.2. Hypomorphic phenotypes may be caused by transcriptional adaptation

Unexpected results were obtained from other knock-in mutants by the Pcry-alleles and fusion- alleles, where DNA constructs were inserted in the third exon. Although encoded proteins from these alleles are expected to be nonfunctional (Fig. S2), their homozygous mutants showed less severe phenotypes than the *Δ*30 homozygous mutants (Table 1-3). Especially, most of the male mutants by the Pcry-alleles showed normal sperm duct development. The results may be caused by a phenomenon called transcriptional adaptation (El-Brolosy et al., 2019; see also Jakutis and Stainier, 2021). Premature termination codons of mutated genes, in many instances, cause degradation of their mRNA through nonsense mediated mRNA decay (see Lykke-Andersen and Jensen, 2015; Nagar et al., 2023). Resulted small RNA fragments induce transcriptional upregulation of genes with similar sequences, which are called adapting genes. Here in the *wnt4a* knock-in mutants, possible candidates of adapting gene are *wnt4b* or other *wnt* genes. Both Pcry- and fusion- alleles contain premature termination codons in the third exon, which may turn on nonsense mediated mRNA decay (Fig. S2).

### 4.3. The function of Wnt4a may be sex and stage dependent

The phenotypes of *Δ*30 homozygous mutants were mostly consistent in both sexes (Table 1). However, that was not the case in other knock-in mutants. The males of the Pcry mutants had normal genital ducts compared to females, about half of which showed failed elongation of the oviducts (Table 3). Similarly, in homozygous mutants of the fusion alleles, most of the females had closed genital duct phenotypes but about half of the males had normal sperm ducts (Table 2). The results may indicate that more Wnt4a activity is required for genital duct formation in the females than in the males. Alternatively, the activities of Wnt4a in the knock-in mutants were higher in the males than in the females. Next, in the knock-in mutants, we found out closure of the genital ducts occurred outside the coelom; the posterior end of the ducts left the coelom but stopped beneath the urinary bladder (Table 2 and 3). These phenotypes were found out in the both sexes and may indicate that Wnt4a signals are required in at least two steps: initial elongation of the duct primordia within the coelom and further posterior elongation beneath the urinary bladder. Again, further cellular and molecular studies on processes of the genital duct elongation are required to answer these questions.

**Table 3.**
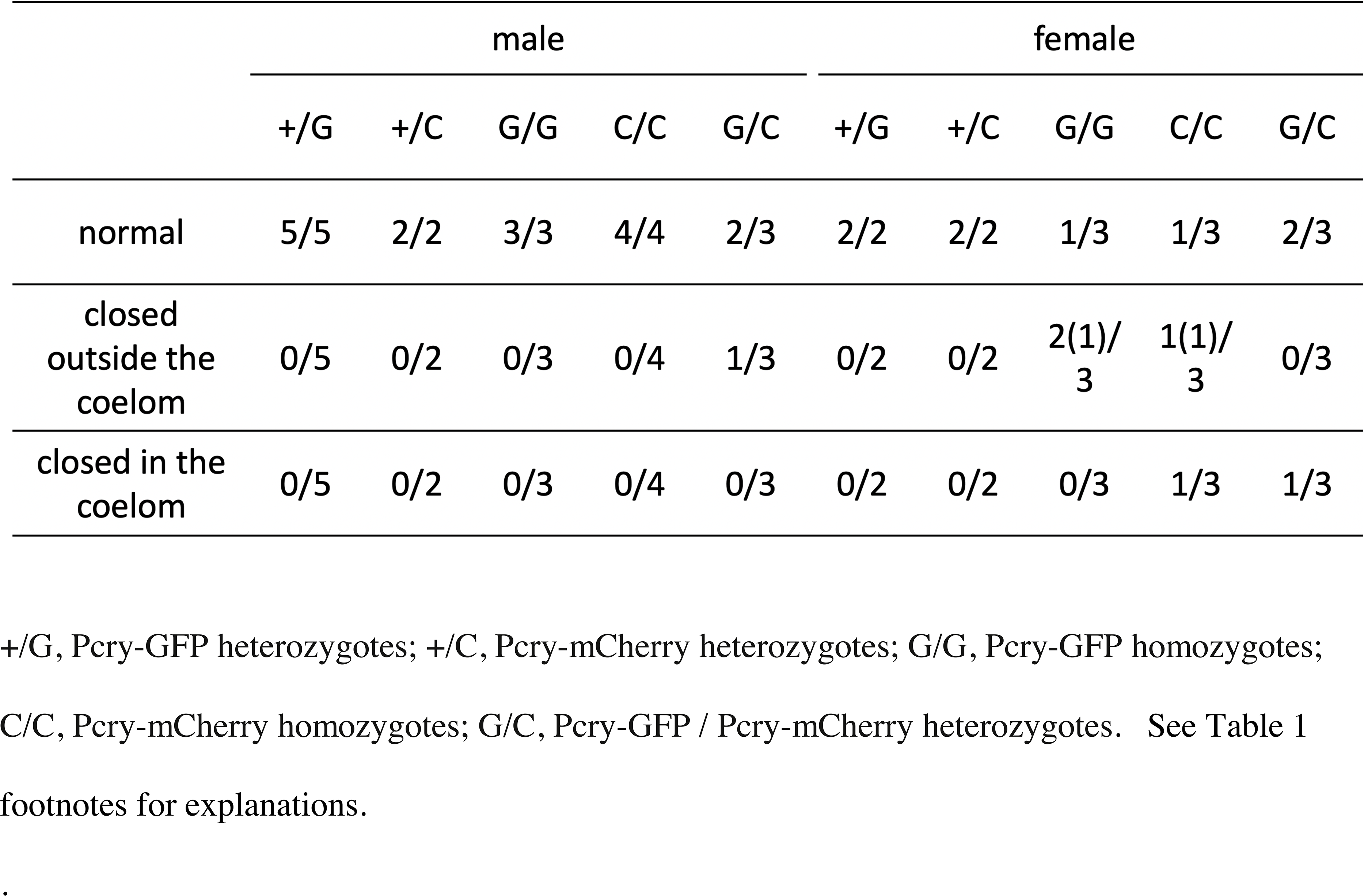
Genital ducts phenotypes of *wnt4a* Pcry allele progenies (G_2_) +/G, Pcry-GFP heterozygotes; +/C, Pcry-mCherry heterozygotes; G/G, Pcry-GFP homozygotes; C/C, Pcry-mCherry homozygotes; G/C, Pcry-GFP / Pcry-mCherry heterozygotes. See Table 1 footnotes for explanations.

**Table 4.**
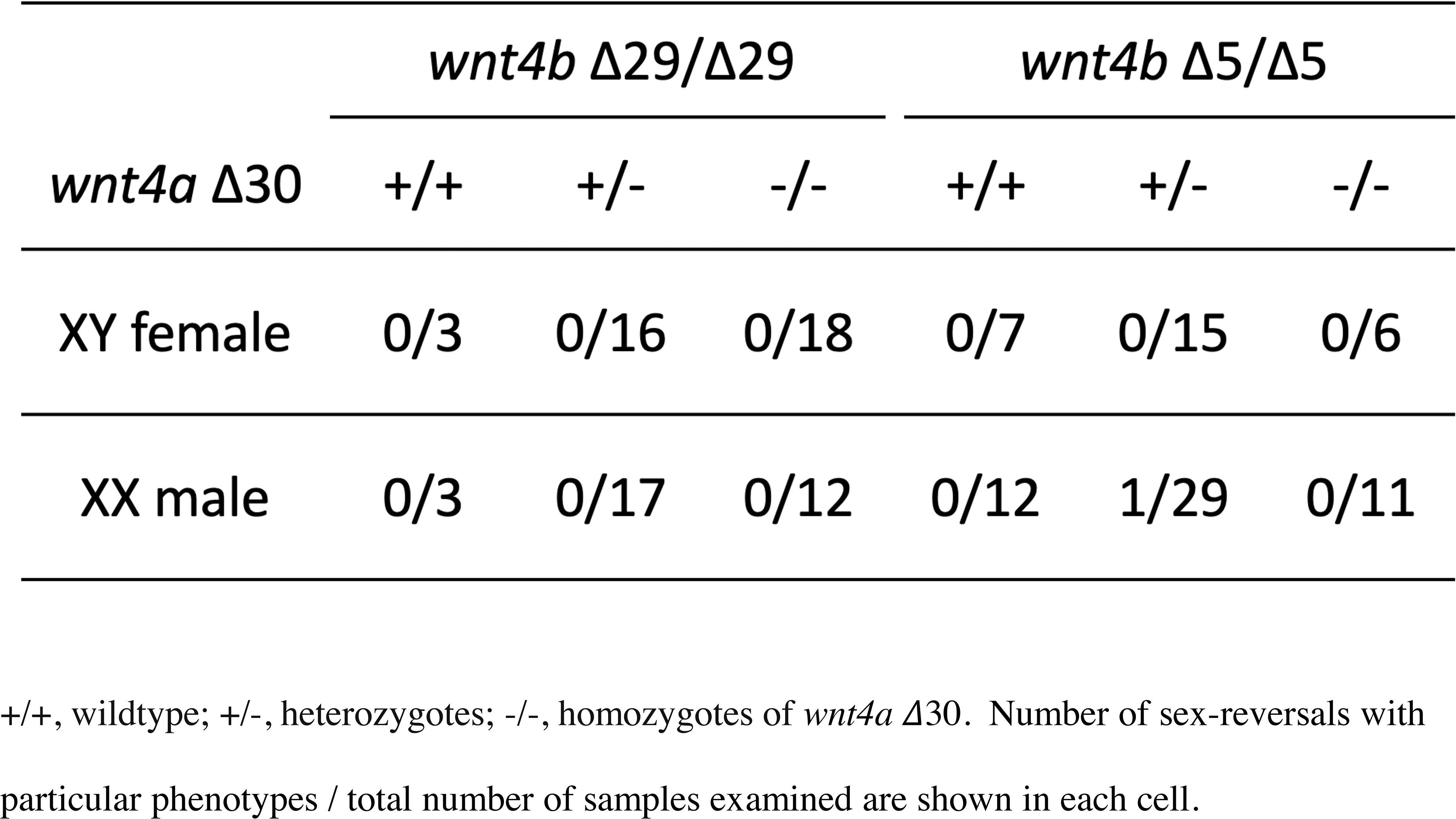
Number of sex-reversals in *wnt4a* / *wnt4b* double mutants +/+, wildtype; +/-, heterozygotes; -/-, homozygotes of *wnt4a Δ*30. Number of sex-reversals with particular phenotypes / total number of samples examined are shown in each cell.

### 4.4. Possible homology between developmental processes of teleost genital ducts and MDs

In mice, *wnt4* KO completely abrogate invagination of the coelomic epithelium and posterior elongation of MDs (Vainio et al., 1999). In teleosts, zebrafish (Kossack et al., 2019) and medaka (the present study), posterior elongation of the genital duct primordia were inhibited in both sexes. In female teleosts, as noted before, the duct epithelium is elongated from epithelium facing the ovarian cavity, which are derived directly from the coelomic epithelium (see Nagahama, 1983; Kanamori et al., 1985). The male epithelium facing the duct lumen are epithelium lining the central canal, which are derived from the Sertoli cells (see Nagahama, 1983; Kanamori et al., 1985). The origin of Sertoli cells in teleosts are lateral plate mesoderm (Nakamura et al., 2006), but there are no conclusive researches showing their origin to be the coelomic epithelium. In mice, however, Sertoli cells are shown to originate from the coelomic epithelium (Karl and Capel, 1999). Similarity in Wnt4 (Wnt4a) mutants in mammals and teleosts strongly suggest presence of homologous processes in genital duct formation in both groups. However, duct shapes and relationships to coelom are very different. In mice, anterior coelomic epithelium are invaginated. In contrast, in teleosts, epithelium within the gonad elongate posteriorly and leave the coelom at the most posterior end (see Fig. 8 for diagrams). Therefore, interpretations as to homology become difficult because of these teleost-specific derived characters. To further argue possible homology, we propose two approaches. One is straightforward molecular and cellular approaches to the detailed studies of medaka genital duct elongation. We may be able to find out common pathways to mice, where detailed studies have been reported (see Mullen and Behringer, 2014). The other approach is that of evo-devo. At least, in sturgeon (Wrobel, 2003), caecillian (amphibian, Wrobel and Süß, 2000), birds (Guioli et al., 2007) and mammals (Guioli et al., 2007; see also Mullen and Behringer, 2014), there are reports indicate that MDs develop as invagination of coelomic epithelium followed by its posterior elongation. The role of Wnt4 (Wnt4a) should be examined in these diverse vertebrate groups.

**Figure 8.**
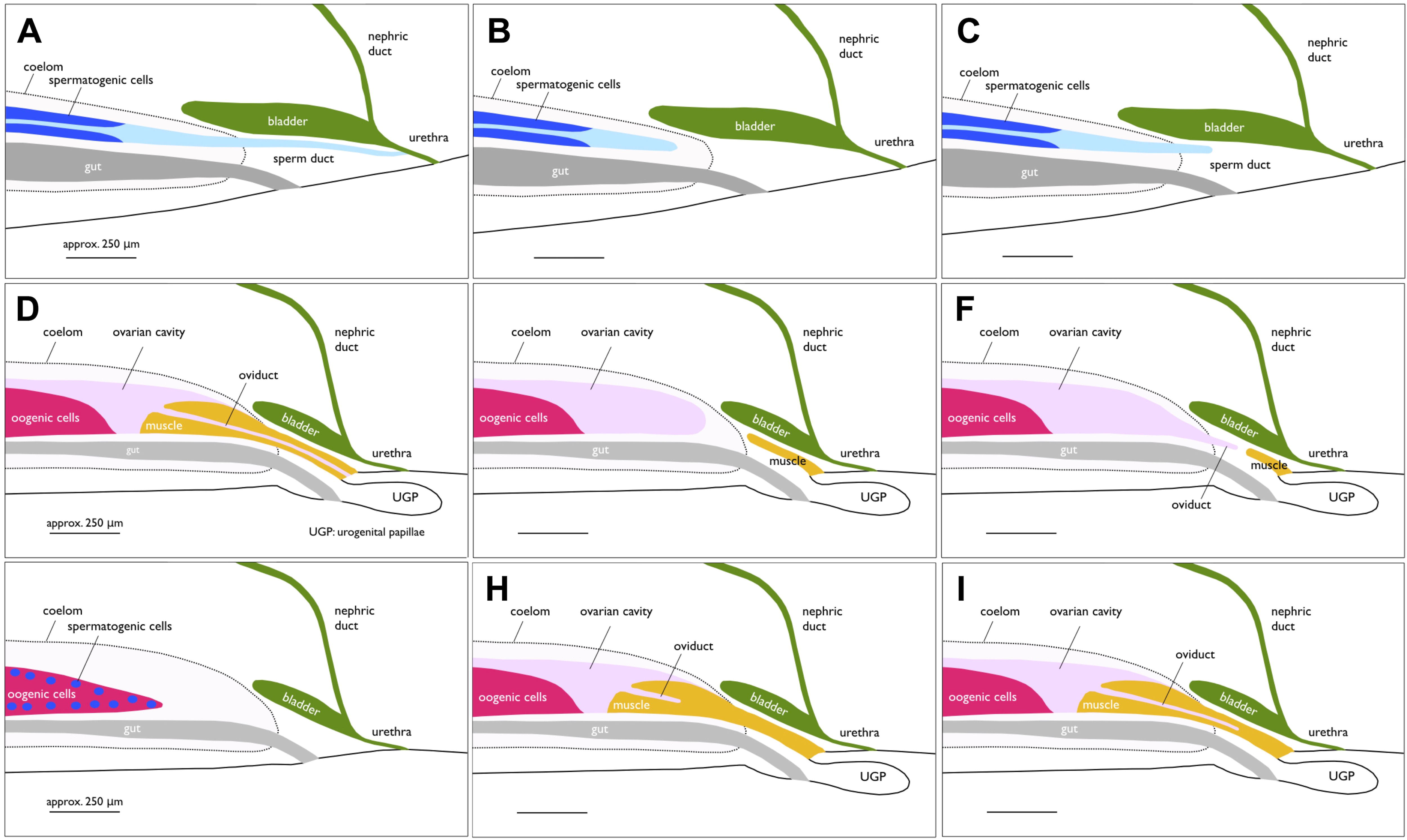
Median plane (sideview) diagrams of urogenital organs in wildtype and *wnt4* mutants. A and D, wildtype male and female, respectively; B and E, mutant male and female with sperm duct closed in the coelom, respectively; C and F, mutant male and female with sperm duct closed outside the coelom, respectively; G, XX *scl* mutant with ovary-like gonad; H and I, female mutants with developed sphincter muscle protruded into the coelom. The oviducts are closed within and outside the coelom, respectively.

**Figure 9.**
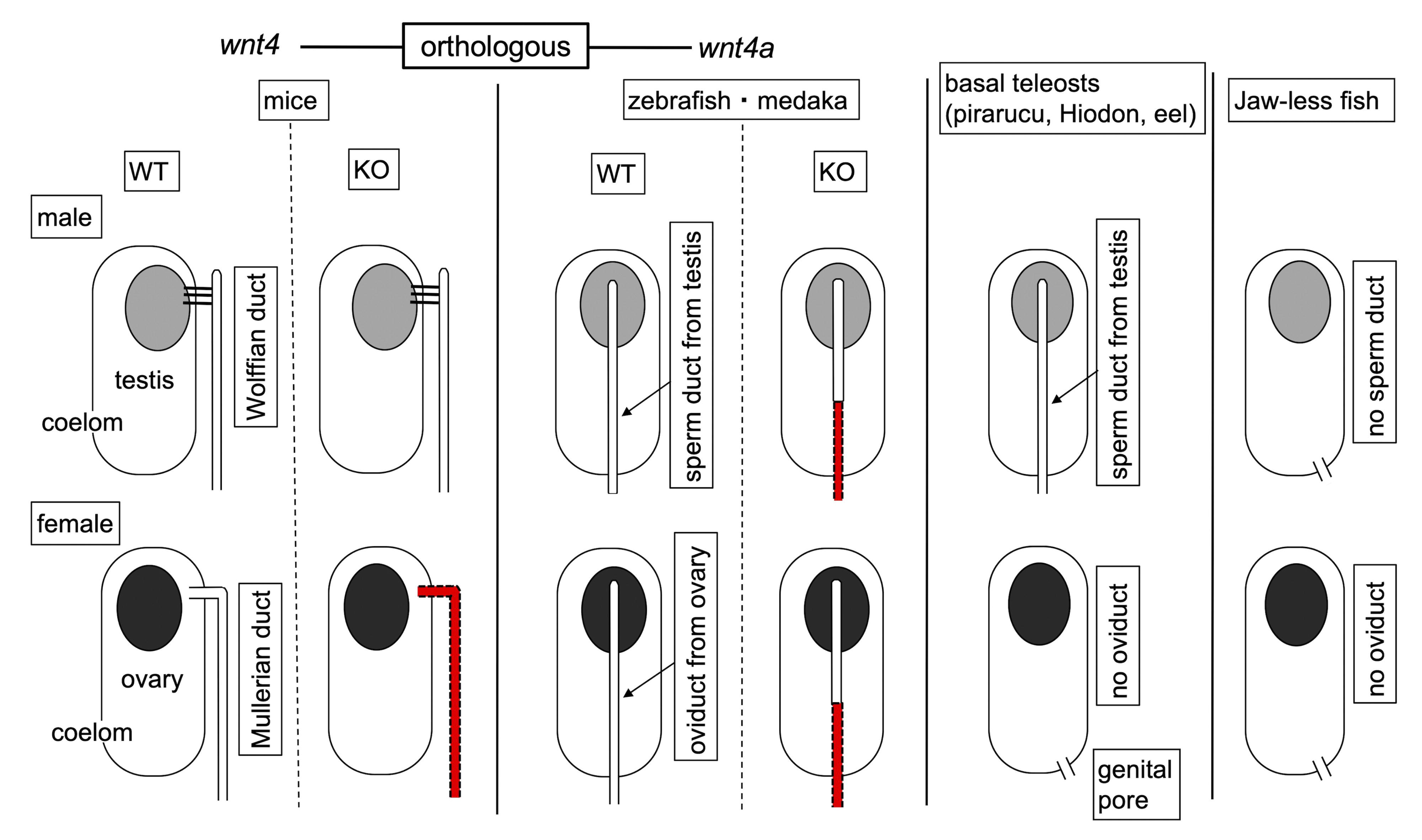
Diagram of genital ducts in various vertebrates and effects of Wnt4 (Wnt4a) mutation (KO). Frontal views with anterior side up. The mutants in orthologous genes, *wnt4* in mice and *wnt4a* in teleosts (zebrafish and medaka), show defective development in MD and genital ducts, respectively (missing tissue shown in red), suggesting presence of homologous developmental processes. Jawless fish of both sexes and females of the basal teleosts lack genital ducts and use genital pores instead, for gamete release.

Cartilaginous fishes (including sharks and rays) first evolved MDs. However, here, pronephric ducts itself differentiate into MDs and WDs develop as longitudinal splitting of the pronephric ducts (see Wourms, 1977; Romer and Parsons, 1977; Goodrich 1930). This striking difference needs to be explained further in cellular and molecular details. Furthermore, jawless fish such as hagfish and lamprey do not have genital ducts; mature sperm or eggs are shed into the coelom and left through genital pores, which are induced by gonadotropins and made by apparent apoptosis of 2-3 layers of cells (Knowles, 1939). These genital pores were argued for or against as MD homologs (see Goodrich, 1930). Finally, basal teleosts such as eels (Tesch, 1977; Fishelson, 1992), Hiodon (Katechis et al., 2007), and pirarucu (Godinho et al., 2005), females do not have genital ducts but have genital pores reminiscent of those of jawless fish. The males do have sperm ducts, although.

At present, there are almost no cellular and molecular descriptions of genital duct or MD development in many vertebrate groups except in mammals. These studies, together with a possible role of Wnt4 (Wnt4a), could shed lights on homology of genital ducts or genital pores among vertebrates.

### 4.5. Gonadal sex differentiation of medaka is not dependent on *wnt4a*

The loss of *wnt4* causes partial female-to-male sex-reversal in mammals (Vainio et al., 1999; see also Bernard and Harley, 2007; Nicol and Yao, 2014). The previous studies revealed that Wnt4 protein acts on gonadal somatic cells to repress transcription of *sox9*, thereby making identity of these cells toward female direction. In teleosts, the ortholog of mammalian *wnt4*, *wnt4a* KO zebrafish showed higher male ratio than the wildtype (Kossack et al., 2019). The authors showed that, in addition to mesenchyme under nephric ducts, *wnt4a* is also expressed in the ovarian somatic cells during gonadal sex differentiation in zebrafish (Kossack et al., 2019). In contrast to zebrafish, we did not detect any effect of *wnt4a* KO on gonadal sex differentiation in medaka. *wnt4a* / *wnt4b* double KO had no effect on the gonadal sex differentiation either. Here, *wnt4a* signals on gonadal somatic cells were not detected in medaka. The difference of wnt4a function in gonadal sex differentiation between medaka and zebrafish may come from genetical sex determination of medaka and environmental sex determination of zebrafish.

In conclusion, we found out medaka *wnt4a* mutants have their genital ducts stopped within the coelom or just after leaving the coelom in both sexes. Together with similar phenotypes reported in zebrafish, we suspect presence of homologous developmental processes between mammals and in teleosts.

## AUTHOR CONTRIBUTIONS

SA, MK, and AO designed and performed CRISPR knockout. TM and TK performed *in situ* hybridization. KK analyzed phylogeny of *wnt4*. NK performed skeletal staining. RK, AO, KH, and AK performed all the other experiments. AK conceived the study and wrote the manuscript.

## DECLARATION OF COMPETING INTEREST

The authors declare that there are no conflicts of interest.

## ACKNOWLEDGMENTS

The authors thanks the following for their help: Dr. H. Hori for genotyping by MultiNA; Dr. H. Hashimoto for *wnt4a* cDNA plasmids; Dr. R. Araki for microCT analyses; National Bioresource Project Medaka (https://shigen.nig.ac.jp/medaka/), especially Dr. M. Matsuda for *scl* mutants. This work was supported in part by a Grant-in-Aid for Scientific Research from JSPS (19K06738 to AK).

## Notes

### Competing Interest Statement

The authors have declared no competing interest.

## REFERENCES

Ansai, S., & Kinoshita, M. (2014) Targeted mutagenesis using CRISPR/Cas system in medaka. Biology Open 3, 362–371.

Bernard, P. & Harley, V.R. (2007) Wnt4 action in gonadal development and sex determination. The International Journal of Biochemistry & Cell Biology 39, 31–43.

Blüm, V. (1986) *Vertebrate Reproduction*. Springer Verlag, Berlin.

El-Brolosy, M.A., Kontarakis, Z., Rossi, A., Kuenne, C., Günther, S., Fukuda, N., Kikhi, K., Boezio, G.L.M., Takacs, C.M., Lai, S.L., Fukuda, R., Gerri, C., Giraldez, A.J., & Stainier D.Y.R. (2019) Genetic compensation triggered by mutant mRNA degradation. Nature 568, 193–197.

Fishelson, L. (1992) Comparative Gonad Morphology and Sexuality of the Muraenidae (Pisces, Teleostei). Copeia 1992, 197–209.

Godinho, H. P., Santos, J. E., Formagio, P. S. & Guimarães-Cruz, R. J. (2005) Gonadal morphology and reproductive traits of the Amazonian fish *Arapaima gigas* (Schinz, 1822). Acta Zoologica (Stockholm) 86, 289–294.

Goodrich, E.S. (1930) Studies on the structure and development of vertebrates. Macmillan, London.

Guioli, S., Sekido, R., & Lovell-Badge, R. (2007) The origin of the Mullerian duct in chick and mouse. Developmental Biology 302, 389–398.

Horie, Y., Myosho, T., Sato, T., Sakaizumi, M., Hamaguchi, S. & Kobayashi, T. (2016) Androgen induces gonadal soma-derived factor, Gsdf, in XX gonads correlated to sex-reversal but not Dmrt1 directly, in the teleost fish, northern medaka (*Oryzias sakaizumii*). Molecular and Cellular Endocrinology 436, 141–149.

Hyodo-Taguchi, Y., & Sakaizumi, M. (1993) List of inbred strains of the medaka, Oryzias latipes, maintained in the Division of Biology, National Institute of Radiological Sciences. Fish Biology Journal MEDAKA 5, 29–30.

Inohaya, K., Takano, Y., & Kudo, A. (2010) Production of Wnt4b by floor plate cells is essential for the segmental patterning of the vertebral column in medaka. Development 137,1807–1813.

Izard, J.W., & Kendall, D.A. (1994) Signal peptides: exquisitely designed transport promoters. Molecular Microbiology 13, 765–73.

Jakutis, G., & Stainier, D.Y.R. (2021) Genotype-Phenotype Relationships in the Context of Transcriptional Adaptation and Genetic Robustness. Annual Review of Genetics 55, 71–91.

Kanamori, A., Nagahama, Y., & Egami, N. (1985) Development of the tissue architecture in the gonads of the medaka *Oryzias latipes*. Zoological Science 2, 695–706.

Karl, J., & Capel, B. (1998) Sertoli cells of the mouse testis originate from the coelomic epithelium. Developmental Biology 203, 23–33.

Katechis, C.T., Sakaris, P.C., & Irwin, E.R. (2007) Population Demographics of *Hiodon tergisus* (Mooneye) in the Lower Tallapoosa River. Southeastern Naturalist 6, 461–470.

Knowles, F.G.W. (1939) The influence of anterior-pituitary and testicular hormones on the sexual maturation of lampreys. Journal of Experimental Biology 16, 535–547.

Kossack, M.E., High, S.K., Hopton, R.E., Yan, Y., Postlethwait, J.H., and Draper, B.W. (2019) Female sex development and reproductive duct formation depend on Wnt4a in zebrafish. Genetics 211, 219–233.

Logan, C.Y. & Nusse, R. (2004) The Wnt signaling pathway in development and disease. Annual Review in Cell Developmental Biology 20, 781–810.

Lombardi, J. (1998) Comparative Vertebrate Reproduction. Kluwer Academic Publishers, Boston.

Loosli, F., Köster, R.W., Carl, M., Kühnlein, R., Henrich, T., Mücke, M., Krone, A., & Wittbrodt, J. (2000) A genetic screen for mutations affecting embryonic development in medaka fish (*Oryzias latipes*). Mechanisms of Development 97, 133–139.

Lykke-Andersen, S., & Jensen, T.H. (2015) Nonsense-Mediated mRNA Decay: An Intricate Machinery That Shapes Transcriptomes. Nature Reviews Molecular Cell Biology 16, 665–677.

Matsuda, M., Nagahama, Y., Shinomiya, A., Sato, T., Matsuda, C., Kobayashi, T., Morrey, C.E., Shibata, N., Asakawa, S., Shimizu, N., Hori, H., Hamaguchi, S., & Sakaizumi, M. (2002) DMY is a Y-specific DM-domain gene required for male development in the medaka fish. Nature 417, 559–563.

Mullen, R.D. & Behringer, R.R. (2014) Molecular Genetics of Müllerian Duct Formation, Regression and Differentiation. Sexual Development 8, 281–296.

Murakami, Y., Ansai, S., Yonemura, A., & Kinoshita, M. (2017) An efficient system for homology- dependent targeted gene integration in medaka (*Oryzias latipes*). Zoological Letters 3, 10.

Nagahama, Y. (1983) The functional morphology of teleost gonads. In Fish Physiology vol. 9 pt. A Ed. by Hoar, W.S., Randall, D.J., Donaldson, E.M. Academic Press, New York, pp 223–264.

Nagar, P., Islam, M.R., & Rahman, M.A. (2023) Nonsense-Mediated mRNA Decay as a Mediator of Tumorigenesis. Genes (Basel) 14, 357.

Nakamura, S., Kobayashi, D., Aoki, Y., Yokoi, H., Ebe, Y., Wittbrodt, J., & Tanaka, M. (2006) Identification and lineage tracing of two populations of somatic gonadal precursors in medaka embryos. Developmental Biology 295, 678–688.

Mehta, S., Hingole, S., & Chaudhary, V. (2021) The Emerging Mechanisms of Wnt Secretion and Signaling in Development. Frontiers in Cell and Developmental Biology 16, 714746.

Nicol, B., & Yao, H.H. (2014) Building an ovary: insights into establishment of somatic cell lineages in the mouse. Sexual Development 8, 243–51.

Owji, H., Nezafata, N., Negahdaripoura, M., Hajiebrahimia, A, & Ghasemia, Y. (2018) A comprehensive review of signal peptides: Structure, roles, and applications. European Journal of Cell Biology 97, 422–441

Romer, A.S., & Parsons, T.S. (1977) The Vertebrate Body, 5th Edition, W. B. Saunders Co., Philadelphia.

Sato, T., Suzuki, A., Shibata, N., Sakaizumi, M., & Hamaguchi, S. (2008) The novel mutant *scl* of the medaka fish, *Oryzias latipes*, shows no secondary sex characters. Zoological Science 25, 299–306

Steinhart, Z., & Angers, S. (2018). Wnt signaling in development and tissue homeostasis. Development 145, dev146589.

Suzuki, A., & Shibata, N. (2004) Process of Genital Ducts in the Medaka, *Oryzias latipes*. Zoological Science 21, 397–406.

Suzuki, A., Tanaka, M., Shibata, N., & Nagahama, Y. (2004) Expression of aromatase mRNA and effects of aromatase inhibitor during ovarian development in the medaka, *Oryzias latipes*. Journal of Experimental Zoology. Part A, Comparative Experimental Biology 301, 266–73.

Tesch, F.W. (1977) The Eel. Chapman and Hall Ltd., New York.

Vainio, S., Heikkilä, M., Kispert, A., Chin, N., & McMahon, A.P. (1999) Female development in mammals is regulated by Wnt-4 signalling. Nature 397, 405–409.

Wourms, J.P. (1977) Reproduction and Development in Chondrichthyan Fishes. American Zoologist 17, 379–410.

Wrobel, K.H. (2003) The genus Acipenser as a model for vertebrate urogenital development: the müllerian duct. Anatomy and Embryology 206, 255–271.

Wrobel, K.H., & Süß, F. (2000) The significance of rudimentary nephrostomial tubules for the origin of the vertebrate gonad. Anatomy and Embryology 201, 273–290.

