## Supplementary material for "Wnt4a is indispensable for genital duct elongation but not for gonadal sex differentiation in the medaka *Oryzias latipes*": Fig. S1-S4; Table S1-3

**Supplementary materials**

Supplementary figures, S1-S4, and tables, S1-S3, are followed. Materials and methods, and References, are included in each legend.

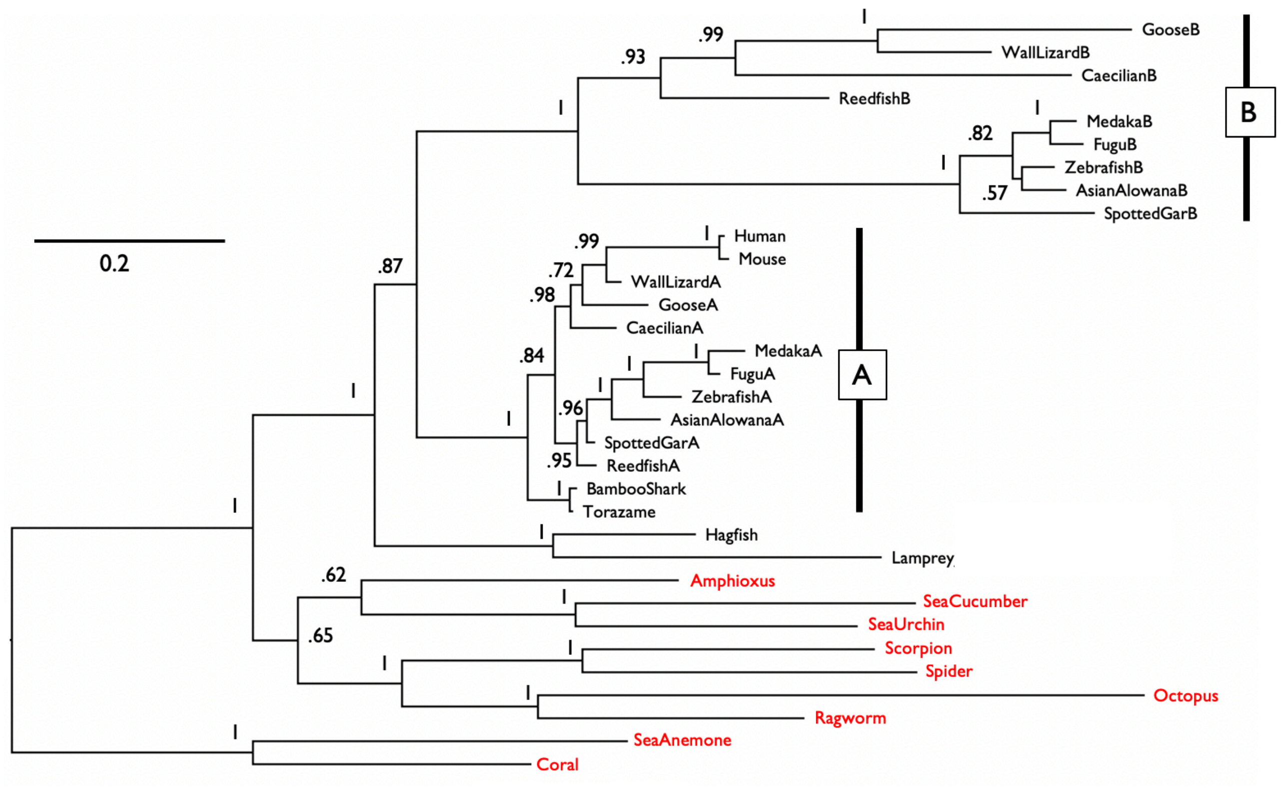


**Figure S1.**

Phylogeny of animal wnt4 proteins. Phylogenetic relationships among species were inferred by maximum-likelihood (ML) and Bayesian inference (BI) approaches. As two methods gave identical tree shape, only BI tree is shown here with posterior probabilities at the nodes. Wnt4 is present in most invertebrates (shown in red) and vertebrates have two wnt4 genes, A and B. Mammalian *wnt4* is orthologous to medaka *wnt4a* and paralogous to medaka *wnt4b*. The accession numbers of the data used are as follows: GooseB, XP_047907497; WallLizardB, XP_028568884; CaecilianB, XP_030043700; ReedfishB, XP_039619571; MedakaB, NP_001153912; FuguB, XP_003966339; ZebrafishB, AAI62132; AsianAlowanaB, XP_018587545; SpottedGarB, XP_006642418; Human, NP_110388; Mouse, NP_033549; WallLizardA, XP_028596829; GooseA, XP_026653200; CaecilianA, XP_030078431; MedakaA, NP_001153911; FuguA, XP_003973796; ZebrafishA, AJG06063; AsianAlowanaA, XP_018614067; SpottedGarA, XP_006641925; ReedfishA, XP_028663537; BambooShark, GCC32830; Torazame, GCB70628; Hagfish, ENSEBUT00000009109.1 (cDNA); Lamprey, XP_032823978; Amphioxus, AAC80431;

SeaCucumber, AMY96432; SeaUrchin, XP_030842454; Scorpion, XP_023222233; Spider, XP_015913740; Octopus, XP_052826593; Ragworm, CAD37166; SeaAnemone, AAV87174; Coral, PFX19278.

Alignment of Wnt4 amino acid sequences was prepared by clustalW (Larkin et al., 2007; https://www.genome.jp/tools-bin/clustalw) with default parameters. The most suitable model of amino acid evolution was selected under the Akaike Information Criterion (Akaike 1974) corrected for small sample sizes (AICc, Hurvich and Tsai 1989) for ML and Bayesian Information Criterion (BIC) (Schwarz 1978) for BI, using ModelTest-NG v. 0.1.7 (Darriba et al. 2020) in ML and Prottest3 (Darriba et al. 2011) in BI. For ML analysis, the JTT-DCMUT+I+G4 model (gamma shape = 0.9025) was determined to be the most appropriate under AICc. The optimal ML tree was constructed using a heuristic search as implemented in RAxML-NG v1.1.0 (Kozlov et al. 2019), with TBR branch swapping. Statistical support for recovered clades was assessed by 10,000 bootstrap replicates (Felsenstein 1985). Bayesian analysis was run with MrBayes 3.2.7a (Ronquist et al. 2012), using the JTT+G model (gamma shape = 0.697) with $5 \times 10^6$ generations Markov chain Monte Carlo (MCMC). Starting trees were random, one cold and three heated chains were run simultaneously. Trees were saved every 1,000 generations. MCMC convergence was confirmed by the average standard deviation of split frequencies (ASDSF) and large effective sample size (ESS >200) in Tracer 1.7.1 (Rambaut et al. 2018). The first 10% of trees were discarded as burn-in and a majority rule (50%) was adopted to determine the posterior probability of clades. The phylogenetic trees were visualized with FigTree ver. 1.4.4 (Rambaut 2023).

### References

Akaike H (1974) A new look at the statistical model identification. IEEE Trans Autom Contr 19:716-723

Darriba D, Posada D, Kozlov AM, Stamatakis A, Morel B, Flouri T (2020) ModelTest-NG: a new and scalable tool for the selection of DNA and protein evolutionary models. Mol Biol Evol 37:291-294

Darriba D, Taboada G, Doallo R, Posada D (2011) ProtTest 3: fast selection of best-fit models of protein evolution. Bioinformatics 27. DOI: https://doi. org/10.1093/bioinformatics/btr088, PMID 21335321

Felsenstein J (1985) Confidence limits on phylogenies: an approach using the bootstrap. Evolution 39:783-791

Hurvich CM, Tsai C-L (1989) Regression and time series model selection in small samples. Biometrika 76:297-307

Kozlov AM, Darriba D, Flouri T, Morel B, Stamatakis A (2019) RAxML-NG: a fast, scalable and user-friendly tool for maximum likelihood phylogenetic inference. Bioinformatics 35:4453-4455

Larkin MA, Blackshields G, Brown NP, Chenna R, McGettigan PA, McWilliam H, Valentin F, Wallace IM, Wilm A, Lopez R, Thompson JD, Gibson TJ, Higgins DG. (2007). Clustal W and Clustal X version 2.0. Bioinformatics, 23, 2947-2948.

Rambaut A (2023) FigTree v1.4.7. Retrieved from https://github.com/rambaut/figtree/releases (accessed April 7, 2023).

Rambaut A, Drummond AJ, Xie D, Baele G, Suchard MA (2018) Posterior summarization in Bayesian phylogenetics using Tracer 1.7. Syst Biol 67:901-904. https://doi.org/910.1093/sysbio/syy1032

Ronquist F, Teslenko M, Van Der Mark P, Ayres DL, Darling A, Höhna S, Larget B, Liu L, Suchard MA, Huelsenbeck JP (2012) MrBayes 3.2: efficient Bayesian phylogenetic inference and model choice across a large model space. Syst Biol 61:539-542

Schwarz G (1978) Estimating the dimension of a model. Ann Stat 6:461-464

A

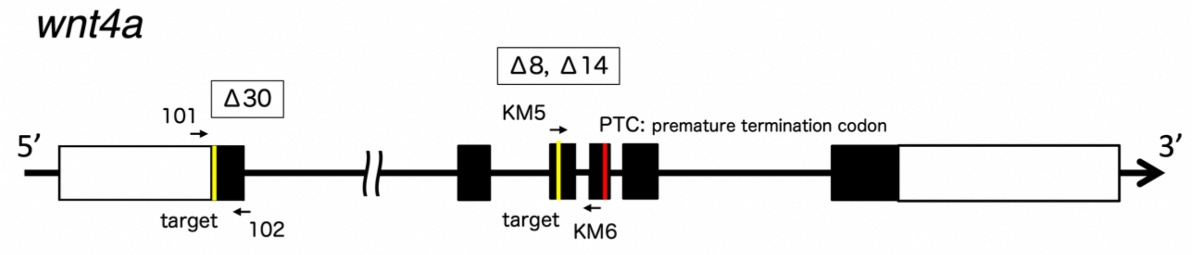


primer sequences for genotyping （5′→3′）

101, GCTCTTGGATTATCCAGAACCC
102, CACTTACAGCCAGTTACTCGCG

PCR products will be 135 bp and 105 bp, respectively for WT and Δ30.

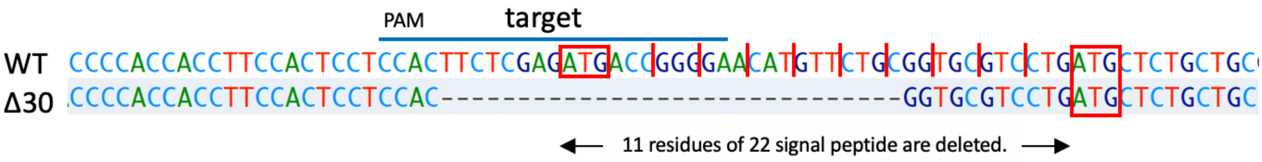


KM5, GTTCAGATCTGTAAGCGTAGCGTG
KM6, GCCTCTCGGGTGCCTAACACC

PCR products will be 250 bp for WT. Genotyping was done by sequencing.

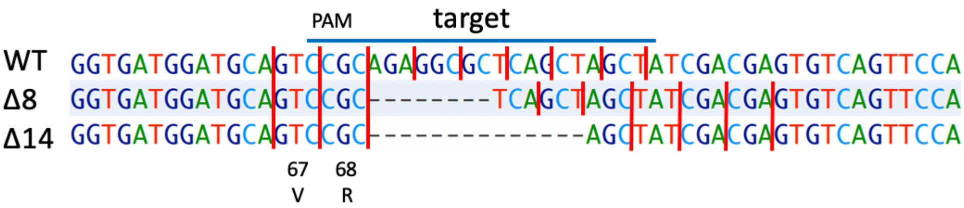


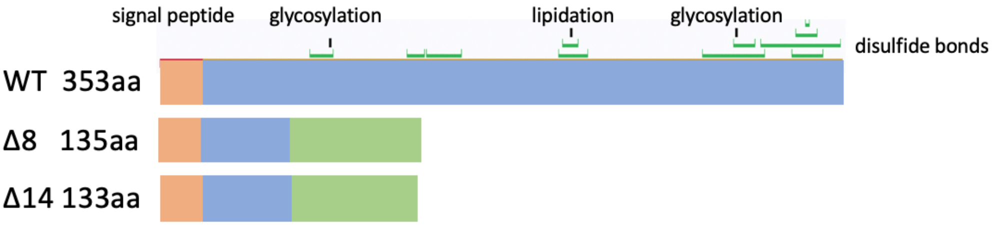

B

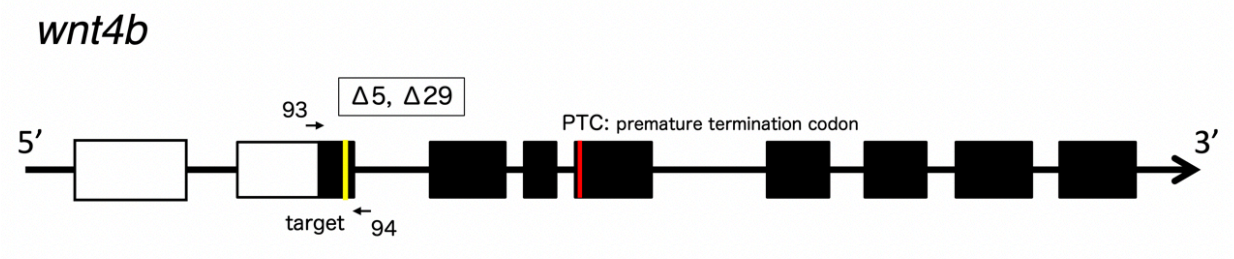


93, AGACAGACCGACAGTCTGATGC
94, CCACGAACACTAGACTTACAGC

PCR products will be 128, 120, 99 bp for WT, Δ8, Δ14, respectively.

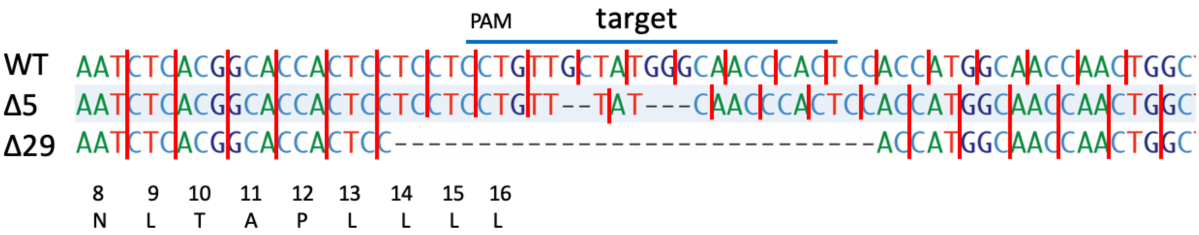


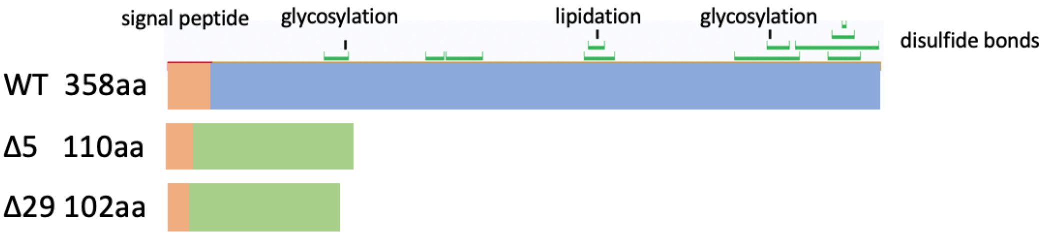

C

Knock-in alleles of *wnt4a*: Transgenes were insertion by CRISPR/cas9 mediated homologous recombination.

1. mouse crystalline promotor-EGFP/mCherry

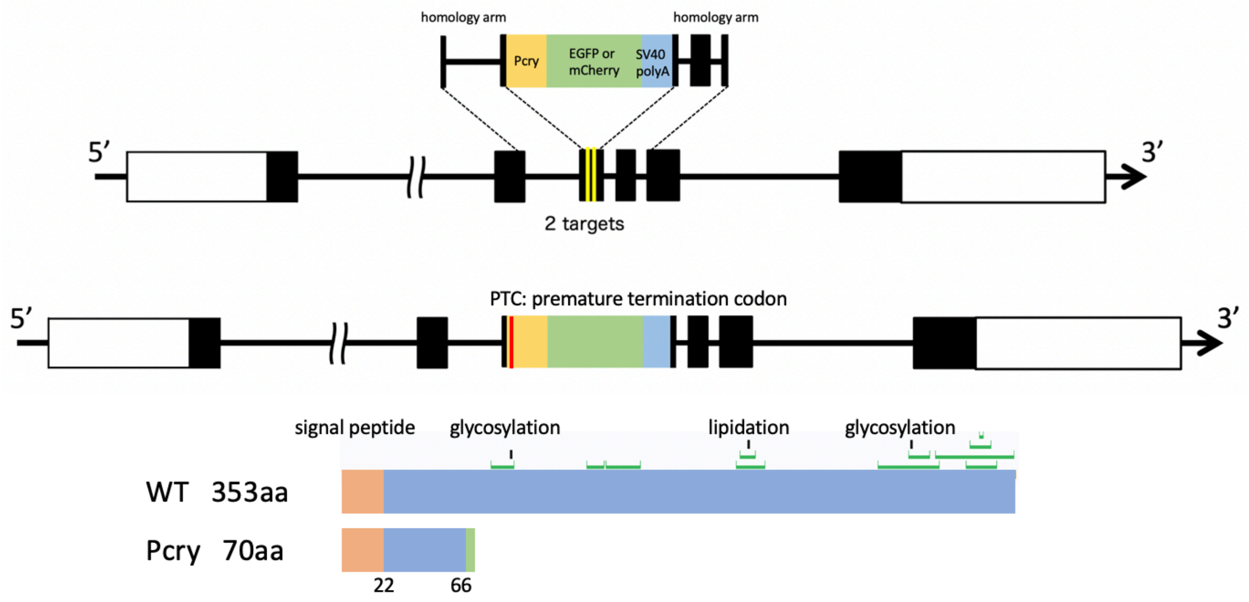


2. WNT4a-EGFP/mCherry fusion

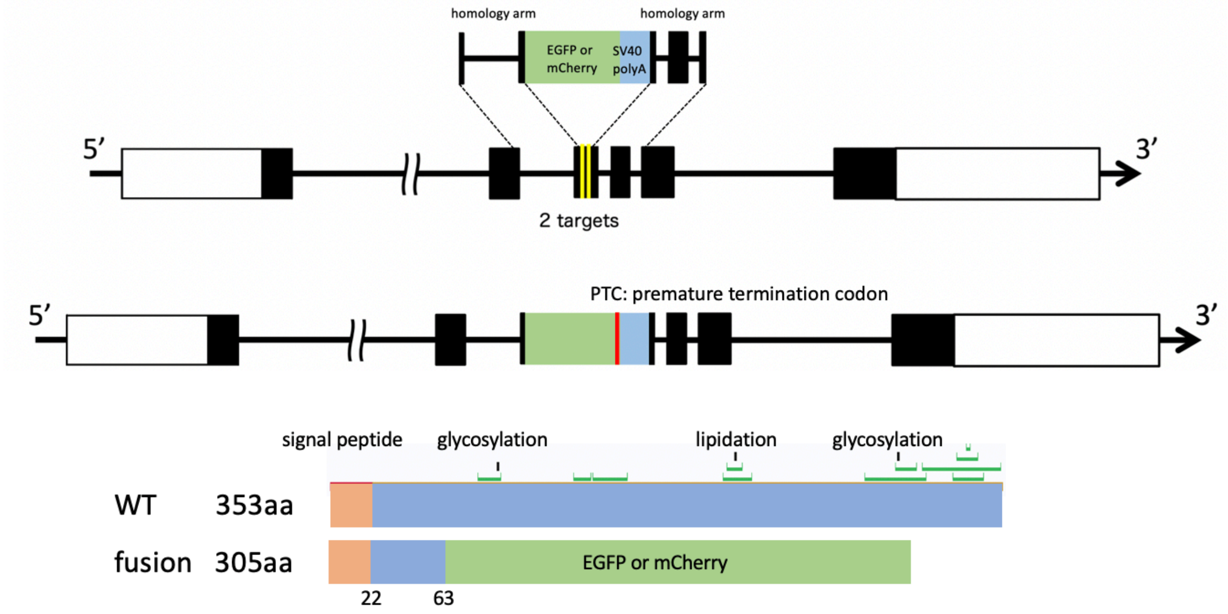


Two target sites are TAGCTAGCTGAGCGCCTCTGCGG and CCGATGGAACTGCTCTACGCTGG.

KM5, GTTCAGATCTGTAAGCGTAGCGTG
KM6, GCCTCTCGGGTGCCTAACACC

PCR products will be 250 bp, 1.1kb, and 1.4 kb for WT, fusion alleles, and Pcry alleles.

D

Genetic sex was determined by following two primer pairs.

1. dmy specific primers (Matsuda et al., 2002)

17.19, 5'-GAACCACAGCTTGAAGACCCCGCTGA-3'
17.20, 5'-GCATCTGCTGGTACTGCTGGTAGTTG-3’

2. dmy/dmrt1 primers

17.z1, 5'-CCGCTGAAAGGCCACAAGCGC-3'
17.z2, 5'-GCCTGCTGCCTCCTCAAGGCG-3'

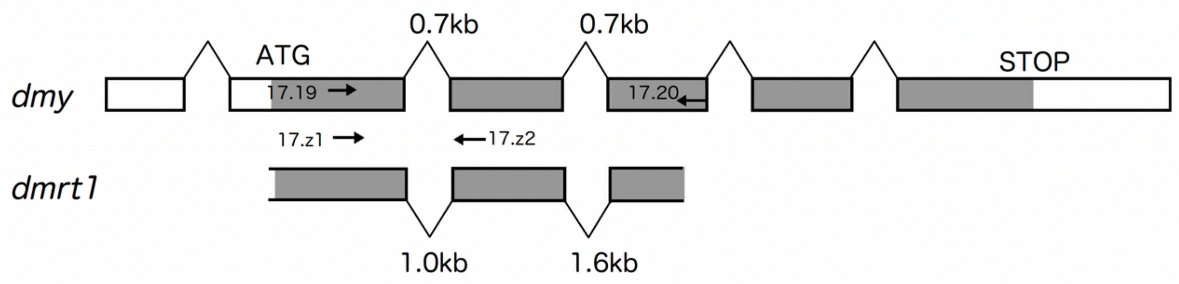


17.19 / 20 : recognize dmy only / target is 1.9 kb
17.z1 / z2 : recognize both *dmy* and *dmrt1* / targets are 0.8 (*dmy*) and 1.1 (*dmrt1*) kb

**Figure S2.**

The mutated alleles of *wnt4a* and *wnt4b* by CRISPR/Cas9. Target sites, mutated sequences, encoded proteins, location of primers for genotyping, and expected PCR products are described. A, deletion alleles of *wnt4a*; B, deletion alleles of *wnt4b*; C, knock-in alleles of *wnt4a*. The exons are shown by rectangles (open, untranslated regions; closed, coding sequence). The functional domains of wnt4 proteins are cited from human WNT4 (UniProt, https://www.uniprot.org/uniprotkb/P56705/entry). In addition, location of two primer pairs used for genetic sex determination and expected PCR products are described in D.

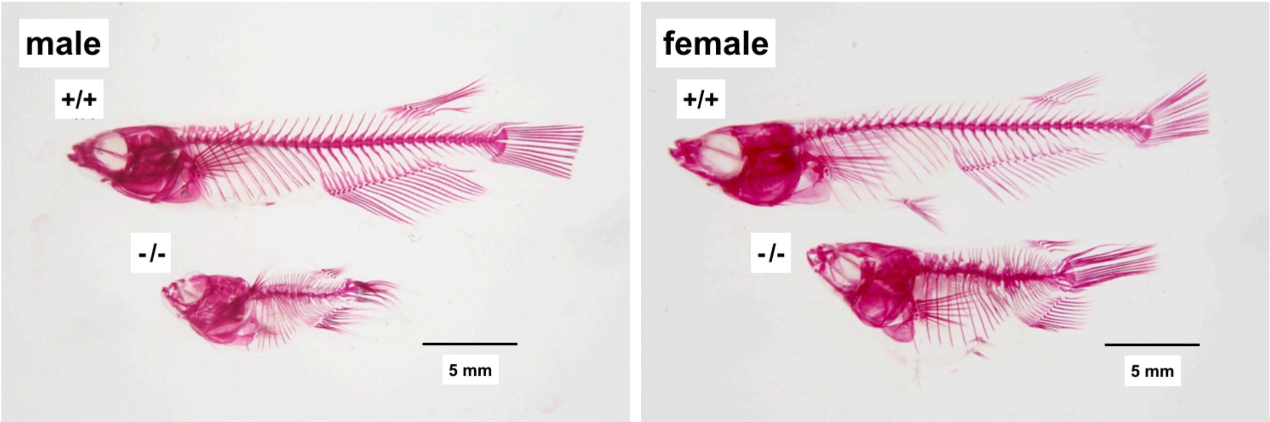


**Figure S3.**

Wildtype (+/+) and *wnt4b Δ*29 mutant (-/-) medaka ($G_2$ from incrossing $G_1$ heterozygotes) were stained with Alizarin Red S according to Dingerkus and Uhler (1977).

Dingerkus, G., & Uhler, L.D. (1977) Enzyme clearing of alcian blue stained whole small vertebrates for demonstration of cartilage. *Stain Technology 52*, 229-32.

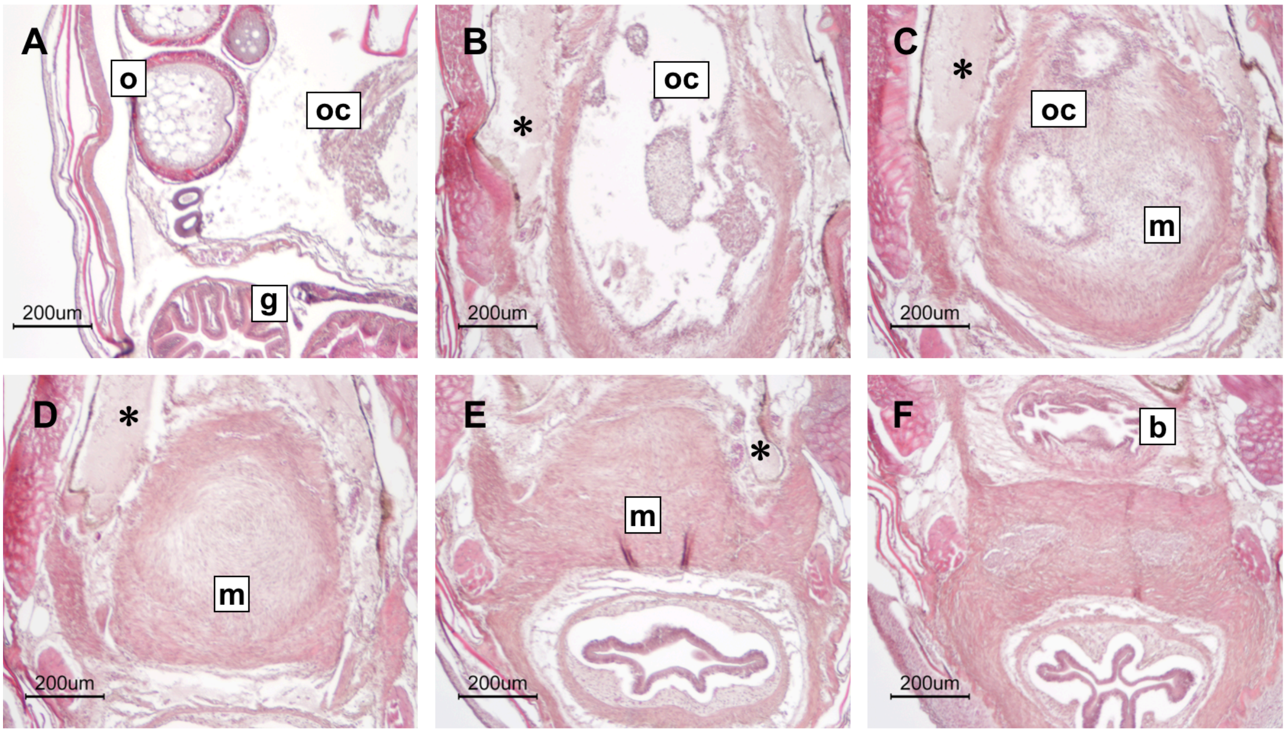


**Figure S4.**

Histology of a female homozygous for *wnt4aΔ*30 ($G_2$ from incrossing $G_1$ heterozygotes) at 4 months after hatching. Cross sections, A to F, are shown in anterior to posterior order. Oogenesis seemed progressing normally (A) and a posterior portion of ovarian cavity (oc) devoid of germ cells (oviduct primordium) was formed (B). In this female, the muscular tissue was well developed and protruded into the coelom and the oviduct primordium stopped elongation within the muscular tissue (C and D). Muscular tissue beneath the urinary bladder and urogenital papillae were developed normally (E and F). o, ovary; g, gut; b, urinary bladder; ugp, urogenital papillae; nd, nephric duct; asterisks, coelom. A sideview diagram is shown in Fig. 8H.

**Table S1.** Genital ducts phenotypes of *wnt4a* Δ8/ Δ14 progenies ($G_1$ and $G_2$ combined)

| | male | | | | female | | | |
|---|---|---|---|---|---|---|---|---|
| | +/+ | +/Δ8 | +/Δ14 | Δ8/Δ14 | +/+ | +/Δ8 | +/Δ14 | Δ8/Δ14 |
| normal | 2/2 | 4/4 | 3/3 | 1/3 | 2/2 | 4/4 | 3/3 | 0/7 |
| closed outside the coelom | 0/2 | 0/4 | 0/3 | 2/3 | 0/2 | 0/4 | 0/3 | 1/7 |
| closed in the coelom | 0/2 | 0/4 | 0/3 | 0/3 | 0/2 | 0/4 | 0/3 | 6(1)/7 |

+/+, wildtype; +/ Δ8, Δ8 heterozygotes; +/ Δ14, Δ14 heterozygotes; Δ8/ Δ14, Δ8/ Δ14 heterozygotes. See Table 1 footnotes for explanations.

**Table S2.** Number of sex-reversals in *wnt4a* mutants (*Δ*30 and fusion alleles)

| | Δ30 | | | fusion | | | |
|---|---|---|---|---|---|---|---|
| | +/+ | +/- | -/- | +/+ | +/G | +/C | G/C |
| XY female | 0/8 | 0/27 | 0/3 | 0/4 | 1/11 | 0/3 | 1/5 |
| XX male | 2/4 | 1/19 | 0/4 | 0/4 | 0/4 | 0/10 | 0/6 |

See Table 4 footnotes for explanations.

**Table S3.** Genital ducts phenotypes of *wnt4a* / *wnt4b* double mutants

| | male | | | | | | female | | | | | |
|---|---|---|---|---|---|---|---|---|---|---|---|---|
| | *wnt4b* Δ29/Δ29 | | | *wnt4b* Δ5/Δ5 | | | *wnt4b* Δ29/Δ29 | | | *wnt4b* Δ5/Δ5 | | |
| *wnt4a* Δ30 | +/+ | +/- | -/- | +/+ | +/- | -/- | +/+ | +/- | -/- | +/+ | +/- | -/- |
| normal | 2/2 | 2/2 | 1/5 | 2/2 | 2/2 | 0/5 | 2/2 | 2/2 | 1/5 | 2/2 | 2/2 | 0/6 |
| closed in the coelom | 0/2 | 0/2 | 4/5 | 0/2 | 0/2 | 5/5 | 0/2 | 0/2 | 4(1)/5 | 0/2 | 0/2 | 6(1)/6 |

+/+, wildtype; +/-, heterozygotes; -/-, homozygotes of *wnt4a* *Δ*30. See Table 1 footnotes for explanations.
